# Unfolded protein response and scaffold independent pheromone MAP kinase signalling control *Verticillium dahliae* growth, development and plant pathogenesis

**DOI:** 10.1101/2020.02.10.941450

**Authors:** Jessica Starke, Rebekka Harting, Isabel Maurus, Rica Bremenkamp, James W. Kronstad, Gerhard H. Braus

**Affiliations:** Department of Molecular Microbiology and Genetics, Institute of Microbiology and Genetics and Göttingen Center for Molecular Biosciences (GZMB), University of Göttingen, Grisebachstr. 8, D-37077 Göttingen, Germany; Department of Microbiology and Immunology, Michael Smith Laboratories, University of British Columbia, Vancouver, BC, V6T 1Z4, Canada

**Keywords:** unfolded protein response, MAPK signalling, MAPK scaffold, oleate Δ12-fatty acid desaturase, *Verticillium dahliae*, plant pathogen

## Abstract

- Development and virulence of the vascular plant pathogen *Verticillium dahliae* are connected and depend on a complex interplay between the unfolded protein response, a Ham5 independent pheromone MAP kinase module and formation of precursors for oxylipin signal molecules.
- Genes coding for the unfolded protein response regulator Hac1, the Ham5 MAPK scaffold protein, and the oleate Δ12-fatty acid desaturase Ode1 were deleted and their functions in growth, differentiation, and virulence on plants were studied using genetic, cell biology, and plant infection experiments.
- The unfolded protein response transcription factor Hac1 is required for initial root colonization, fungal conidiation and propagation inside the host and is essential for resting structure formation. Microsclerotia development, growth and virulence require the pheromone response MAPK pathway, but without the Ham5 scaffold function. Single ER-associated enzymes for linoleic acid production make important contributions to fungal growth but have only a minor impact on the pathogenicity of *V. dahliae*.
- Fungal growth, sporulation, dormant structure formation and plant infection require a network of the Hac1-regulated unfolded protein response, a scaffold-independent pheromone response MAPK pathway and formation of precursors for signalling. This network includes interesting targets for disease management of the vascular pathogen *V. dahliae*.

## Introduction

*Verticillium dahliae* is a soil-borne asexual ascomycete causing vascular wilting disease in a broad range of plants including high value crops (Pegg & Brady, 2002; Luo *et al*., 2014). Unfavourable conditions induce the formation of highly resistant, black melanized microsclerotia as characteristic dormant structures which persist in the soil and can overwinter for at least 14 years (Wilhelm, 1955; Griffiths, 1970; Pegg & Brady, 2002). The fungus germinates upon recognition of an appropriate host and enters the plant preferably via natural root wounds, root tips or lateral root hairs (Fitzell *et al*., 1980; Eynck *et al*., 2007; Vallad & Subbarao, 2008; Su *et al*., 2018). Hyphae grow from cortical cells towards the central cylinder and some of them successfully reach the xylem (Klosterman *et al*., 2009). Asexual spores are formed and spread within the vascular system via the transpiration stream (Klosterman *et al*., 2009). Colonization of tissues neighbouring the xylem correlates with induction of disease symptoms (Fradin & Thomma, 2006). Limited nutrient availability in the dying host or in plant debris induces the formation of microsclerotia for persistence (Fradin & Thomma, 2006). During the infection cycle signals from the host environment induce differentiated fungal development and activate specific responses to enable colonization and suppression of the plants immune system.

The unfolded protein response (UPR) pathway monitors protein folding and secretion capacities in the endoplasmic reticulum (ER) lumen and mediates expression of genes involved in ER stress relief (Kozutsumi *et al*., 1988; Kohno *et al*., 1993; Hetz, 2012; Heimel, 2015). Sensing of un- or misfolded proteins in the ER lumen depends on transmembrane sensors as Ire1 described in yeast (Mori *et al*., 1993; Okamura *et al*., 2000). The cytoplasmic endoribonuclease domain of Ire1 is responsible for unconventional splicing of the mRNA encoding the bZIP transcription factor Hac1 (Sidrauski & Walter, 1997; Gonzalez *et al*., 1999). Splicing of the uninduced *HAC1* (*HAC1^u^*) mRNA results in the induced *HAC1* (*HAC1^i^*) mRNA variant. The *HAC1^i^* mRNA is translated into the Hac1 protein which regulates UPR target genes, encoding, for example, chaperones, glycosylation enzymes, proteins required for vesicle transport, lipid biosynthesis or regulators for adaptation of ER size (Cox *et al*., 1993; Mori *et al*., 1996, 1998; Kaufman, 1999; Travers *et al*., 2000; Conn, 2011; Hetz, 2012).

The role of the UPR transcription factor Hac1 homologs and orthologues varies within human or plant pathogenic fungal species (Krishnan & Askew, 2014). The ER stress response mechanism of the opportunistic human pathogenic yeast *Candida glabrata* is regulated in an Ire1-dependent decay independently of Hac1 (Kimmig *et al*., 2012; Miyazaki *et al*., 2013). This results in splicing of various ER-associated mRNAs by Ire1 to cope with ER stress (Heimel *et al*., 2013; Miyazaki *et al*., 2013). Necrotrophic plant ascomycetes such as the rice blast fungus *Magnaporthe oryzae* require Hac1 and the UPR for conidia formation as well as for penetration and invasive hyphae growth during plant infection (Tang *et al*., 2015), whereas in *Alternaria brassicicola* the UPR is required for resistance against plant antimicrobial compounds and virulence, but is not involved in plant penetration and initial colonization (Joubert *et al*., 2011; Guillemette *et al*., 2014). The UPR pathway regulates virulence specific genes in the dimorphic basidiomycete corn smut fungus *Ustilago maydis* (Heimel *et al*., 2010; Hampel *et al*., 2016; Pinter *et al*., 2019). In this fungus a cross-regulation of the UPR pathway and the pheromone response mitogen-activated protein kinase (MAPK) pathway was described. Pathogenicity of the smut fungus depends on activation of the UPR after invasion of the plant surface (Heimel *et al*., 2010, 2013). When activated prior to entering the plant cell, the UPR inhibits filamentous growth and virulence by reduction of the pheromone response MAPK pathway activity through Rok1-dependent dephosphorylation of the MAPK Kpp2 (Schmitz *et al*., 2019).

The pheromone response MAPK pathway was originally described in *Saccharomyces cerevisiae* for a-cell-α-cell fusion during mating where the core module consists of the Ste50 adaptor, the Ste11 MAP kinase kinase kinase (MAP3K), the Ste7 MAP kinase kinase (MAP2K), and the Fus3 MAP kinase (MAPK). A phosphorylation signal is sequentially transferred with the help of the scaffold protein Ste5 to regulate downstream targets such as the transcription factor Ste12 (Alvaro & Thorner, 2016). Components of this pheromone MAPK core can also participate in other Ste5 independent MAPK pathways, such as the starvation mediated filamentous growth pathway of yeast comprised of the Ste50 adaptor, the MAP3K Ste11 and MAP2K Ste7, and the MAPK Kss1 (Cullen *et al*., 2012). The pheromone response MAPK pathway plays an important role in the pathogenicity of numerous filamentous fungi (Jiang *et al*., 2018).

The *V. dahliae* pheromone response MAPK pathway is essential for pathogenicity and includes the upstream adaptor Vste50, Ste11 (MAP3K), Mek2 (Vst7; MAP2K) corresponding to yeast Ste7, Vmk1 (MAPK) corresponding to Fus3 and the downstream Ste12-like transcription factor Vph1 (Rauyaree *et al*., 2005; Qi *et al*., 2016; Sarmiento-Villamil *et al*., 2018; Li *et al*., 2019; Yu *et al*., 2019). Vmk1, Mek2, Ste11, and Vste50-deficient mutant display defects in microsclerotia development, whereas *VPH1* deletion strains are unaffected. Oxygenated polyunsaturated fatty acids (oxylipins) are fungal hormones, which modulate fungal development, pathogenicity and mycotoxin production (Fischer & Keller, 2016). Oxylipins can trigger the biosynthesis of lipid metabolites with functions in plant host colonization, and can manipulate the host lipid metabolism and alter plant defence responses presumably by mimicking endogenous signal molecules. The interplay between plant and fungal oxylipins can direct the outcome of the fungus-plant interaction (Brodhun & Feussner, 2011). The major precursor of fungal oxylipins is the polyunsaturated fatty acid linoleic acid (18:2Δ9,12). Linoleic acid is synthesized by oleate Δ12-fatty acid desaturases such as OdeA from oleic acid (18:1Δ9) by introduction of a second double bond into the carbon chain at position 12 from the carboxy-terminus (Uttaro, 2006). OdeA is required for asexual and sexual development as well as the formation of resting structures in different Aspergilli (Calvo *et al*., 2001; Chang *et al*., 2004; Wilson *et al*., 2004).

In this study, we analysed the impact of the unfolded protein response and the pheromone response MAPK pathway on the development and pathogenicity of the vascular plant pathogen *V. dahliae.* We show that a functional UPR is required for successful colonization of its host. The UPR regulator Hac1 is essential for the formation of dormant structures, which is critical for the ability of the fungus to persist in the soil for many years and to successfully re-establish the disease in the next season. A functional UPR also contributes to the growth and conidiation of *V. dahliae*. The UPR might crosstalk with the pheromone response MAPK pathway, which does not require the Ham5 scaffold protein (Dettmann *et al*., 2014; Jonkers *et al*., 2014; Frawley *et al*., 2018) to promote *V. dahliae* resting structure formation, growth and virulence. The ER-associated oleate Δ12-fatty acid desaturase producing linoleic acid as precursor of oxylipin hormones primarily promotes fungal growth with only minor impacts on pathogenicity of *V. dahliae*. This analysis suggests an intricate interplay between UPR and pheromone MAPK signalling for fungal growth, development and virulence in the vascular pathogen *V. dahliae*.

## Material and Methods

*V. dahliae* JR2 (Fradin *et al*., 2009) was used as wildtype for construction of all Verticillium strains used in this study. Southern hybridizations of constructed strains are shown in Supporting Information Figures S1, S2 and S3. All Verticillium strains are listed in Supporting Information Table S1, primers in Table S2 and plasmids in Table S3. Methods for bacterial and fungal cultivation and strain construction are described in Supporting Information Methods S1.

### Quantification of growth and developmental structures

Verticillium growth was quantified as colony diameter over time. Melanization of colony centres was quantified by determination of the brightness factor using ImageJ software. Conidia were quantified from liquid SXM cultures. For details see Supporting Information Methods S1.

### Confocal microscopy

Localization of Ode1-GFP was examined by confocal fluorescence microscopy by inoculation of ∼50 000-100 000 freshly harvested spores in 300 µl liquid PDM per well in µ-slide 8 well microscopy chambers (ibidi) and incubation at 25 °C for the indicated time. Hyphal morphology and subcellular localization were observed with a Plan-Neofluar 100x/1.4 oil objective (Zeiss, 300 ms exposure time for GFP signals, 800 ms exposure time for RFP signals). Vacuoles were visualized by fluorescence microscopy after staining with 0.3 µl FM4-64 Dye (Thermo Fisher Scientific) for one hour.

### Protein and nucleic acid purification and hybridization

RNA was purified using the Direct-zol RNA MiniPrep Kit. Reverse transcription of RNA was performed using the QuantiTect Reverse Transcription Kit (Qiagen). Genomic DNA isolation, Southern hybridization, protein extraction and immunoblots were performed according to standard protocols. For details see Supporting Information Methods S1.

### Quantification of gene expression

Expression levels of *HAC1* were quantified relative to the reference genes histone *H2A* (*VDAG_JR2_Chr4g01430a*) and *EIF2B (VDAG_JR2_Chr4g00410a*) by qRT PCR. For details see Supporting Information Methods S1.

### *Arabidopsis thaliana* root colonization assay

The root colonization assay was modified from Tran *et al*., 2014. Three-week-old *A. thaliana* (Col-0) plants were infected with conidial suspensions (1×10^5^ spores/ml) and fluorescence microscopy of the roots was conducted at the indicated time points. For details see Supporting Information Methods S1.

### Plant infection experiments

Ten-day-old seedlings of *Solanum lycopersicum* (Moneymaker) were infected by root dipping into a solution of 1×10^7^ conidiospores/ml and disease symptoms were scored after 21 d. Discoloration of the hypocotyl was determined by observation of cross sections. To test for fungal outgrowth from infected plants, stems were harvested, surface sterilized and incubated on PDM plates supplemented with chloramphenicol (100 µg/ml) at 25 °C for 7 d. For details see Supporting Information Methods S1.

### Sequence analyses

The online databases National Center for Biotechnology Information (NCBI; Geer *et al*., 2010) and Ensembl Fungi (Kersey *et al*., 2018) were used for BLAST searches. Verticillium gene predictions, accession numbers, and sequences were obtained from Ensembl Fungi. The web server RNAfold (Zuker, 2003) was used for determination of secondary structures in mRNAs. Prediction of protein domains was performed using the InterPro website (http://www.ebi.ac.uk/Tools/pfa/iprscan; Jones *et al*., 2014). Presence of nuclear localization signals was analysed using the online databases cNLS mapper (http://nls-mapper.iab.keio.ac.jp/cgi-bin/NLS_Mapper_form.cgi Kosugi *et al*., 2009), and DeepLoc-1.0 (Almagro Armenteros *et al*., 2017; http://www.cbs.dtu.dk/services/DeepLoc/index.php). Multiple alignment of protein sequences was performed with ClustalW (Thompson *et al*., 1994) or Muscle (Edgar, 2004) algorithms in MEGA6.0 software (Tamura *et al*., 2013). Phylogenetic analysis was conducted using Maximum likelihood tree calculations with MEGA6.0 software.

## Results

### The *V. dahliae* unfolded protein response pathway regulator Hac1 supports fungal growth and is essential for resting structure development

The function of the UPR pathway regulator Hac1 for growth, development or virulence has not yet been analysed in fungal plant vascular pathogens. The UPR is important because it monitors secretion capacity and ER protein folding. In particular, the controlled formation of secreted effector proteins is critical for fungal differentiation processes and colonization-related adaptation to provide appropriate responses to different environmental signals, which are perceived at the cell membrane, transduced to the nucleus and then change ER mediated secretion.

A *HAC1* homolog encoding the UPR transcription factor was identified in *V. dahliae* by BLAST search. The *HAC1* gene carries a 53 nucleotides (nt) conventional intron and an additional non-conventional 20 nt intron. Splicing of the conventional intron results in a *HAC1^u^* transcript of 1581 nt encoding a protein with 526 aa. Additional splicing of the non-conventional intron results in *HAC1^i^* with an altered reading frame of 1254 nt encoding for the 417 aa Hac1 protein. Expression of the larger *HAC1^u^* as well as the smaller *HAC1^i^* splice variants were verified by characterizing amplified cDNA sequences. Only *HAC1^i^* transcripts encoding Hac1 were found in the presence of ER stress mediated by DTT (induced conditions), whereas without DTT (uninduced conditions) both *HAC1^u^* and smaller *HAC1^i^* were identified (Fig. 1a-c, Supporting Information Table S4).

**Figure 1:**
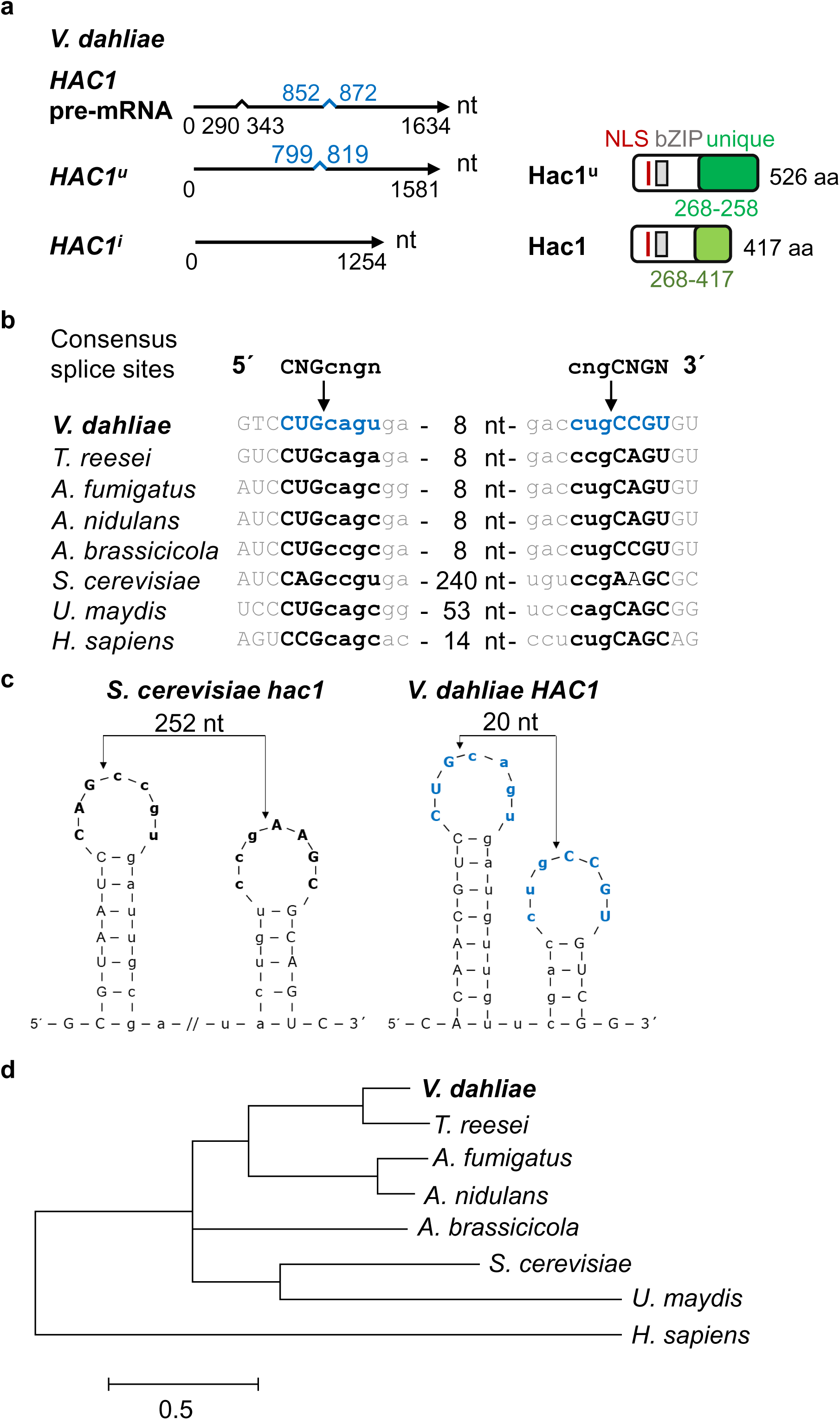
The *V. dahliae HAC1* gene of the unfolded protein response (UPR). (**a**) *Verticillium dahliae HAC1* mRNA and protein variants. Unspliced *HAC1* pre-mRNA (1634 nt) contains a conventional (53 nt, black) and an unconventional (20 nt, blue) intron. *HAC1* produces two mRNA splice variants: Splicing of only the conventional intron results in *HAC1^u^* mRNA (1581 nt) for the potential Hac1^u^ protein (526 aa); additional unconventional splicing results in the shorter induced *HAC1^i^* ORF (1254 nt), encoding the 417 aa Hac1 protein. Both proteins possess identical N-termini (268 aa) with basic leucine zipper domain (bZIP, grey, PS50217; 107-164 aa), and nuclear localization signal (NLS, red, 94 to 105 aa), whereas Hac1^u^ and Hac1 C-termini are unique (dark vs. light green). (**b**) Alignment of 5’ and 3’ splice sequences of unconventionally spliced introns shows high conservation of *V. dahliae HAC1, Trichoderma reesei hac1* (M419DRAFT_128619), *Aspergillus fumigatus hacA* (XM_743634), *Aspergillus nidulans hacA* (AN9397), *Alternaria brassicicola HacA* (Joubert *et al*., 2011), *Sacharomyces cerevisiae hac1* (NC_001138.5), *Ustilago maydis cib1 (UMAG_11782)* and *Homo sapiens XBP1* (NM_005080.3) (CNG’CNGN = consensus sequence (Hooks & Griffiths-Jones, 2011), arrows = cleavage sites, lowercase characters = intron sequences, capital letters = splice sequences, numbers of nucleotides are given for those not shown). (**c**) Predicted twin stem-loop secondary structures of 5’ and 3’ splice sequences of unconventional *V. dahliae* and *S. cerevisiae HAC1* introns. (Arrows = cleavage sites, bold characters = splice sequences, lowercase characters = introns, // = discontinuation of intron sequence) (**d**) Phylogenetic tree derived of Hac1-like proteins with *Verticillium dahliae* Hac1 (Table S5), *Trichoderma reesei* HACI^i^ (XP_006964054.1), *Aspergillus fumigatus* HacA^i^ (ACJ61678.1), *Aspergillus nidulans* HacA^i^ (Q8TFU8.2), *Alternaria brassicicola* AbHacA (Joubert *et al*., 2011), *Saccharomyces cerevisiae* HAC1^i^ (NP_116622.1), *Ustilago maydis* cib1^s^ (XP_011390112.1), and *Homo sapiens* XBP1 (NP_001073007.1) sequences (Muscle algorithm, scale bar = average number of amino acid substitutions per site).

The length of 20 nt for the unconventionally spliced intron of *V. dahliae HAC1* is similar to other filamentous ascomycetes (20-26 nt), but smaller than in the dimorphic basidiomycete *U. maydis* (65 nt) or in budding yeast (252 nt) (Fig. 1b). Unconventional splicing of mRNAs of *HAC1* homologs requires the cytosolic endoribonuclease domain of the ER membrane resident sensor Ire1. This domain recognizes the conserved consensus splice sites 5’-CNG’CNGN-3’ (Hooks & Griffiths-Jones, 2011a). 5’ and 3’ intron-exon-borders of the 20 nt intron from *V. dahliae HAC1* are conserved (Fig. 1b). Similar to Hac1 splice sites of different organisms, the consensus splice site of the unconventional *V. dahliae HAC1* intron was predicted to form a characteristic twin stem-loop secondary structure (Fig. 1c).

The two *HAC1* splice variants encode proteins with identical N-but different C-terminal regions. The shared N-terminal 268 aa region includes the NLS motif and the bZIP domain (Fig. 1a). The remaining C-termini of the deduced larger 526 aa Hac1^u^ protein of 58 kDa and the smaller 417 aa Hac1 protein of 44 kDa are unique (Fig. 1a, the amino acid sequence of Hac1 is shown in Table S5). A phylogenetic analysis revealed similarities between the *V. dahliae* Hac1 protein encoded by the unconventionally spliced mRNA *HAC1^i^* to described UPR regulatory proteins in other fungi and to human XBP1 (X-box binding protein) (Fig. 1d). Proteins from *V. dahliae* and *T. reesei* cluster in one subclade. The Hac1 protein of *S. cerevisiae* showed higher similarity to the *U. maydis* Cib1 protein than to *V. dahliae* Hac1.

A deletion strain was constructed to analyse the role of *HAC1* in the development and virulence of *V. dahliae.* This mutant strain was used for ectopic reintegration of either the entire *HAC1* gene (*HAC1-C*) or one of the two mRNA *HAC1* variants (Fig. 1a) fused to *HA* at the 3’-end and under control of the native promoter and terminator. Expression levels of induced and uninduced *HAC1* mRNA variants were examined with the same primer pair in the constructed strains in comparison to wildtype and Δ*HAC1*. The *HAC1-C* complementation strain displayed wildtype-like *HAC1* expression levels, whereas expression was reduced to less than 40% in *HAC1^u^-HA*, and even to ∼60% in *HAC1^i^-HA* (Fig. 2a). Immunoblot analysis was performed to investigate whether both *HAC1* mRNA variants were translated into Hac1^u^-HA and Hac1-HA fusion proteins. A band at ∼70 kDa instead of the predicted 46 kDa was obtained for *HAC1^i^*-*HA,* expressing the unconventionally spliced *HAC1* mRNA variant, whereas there were no signals for the *HAC1^u^-HA* strain containing the mRNA variant where only the conventional intron was spliced (Fig. 2b).

**Figure 2:**
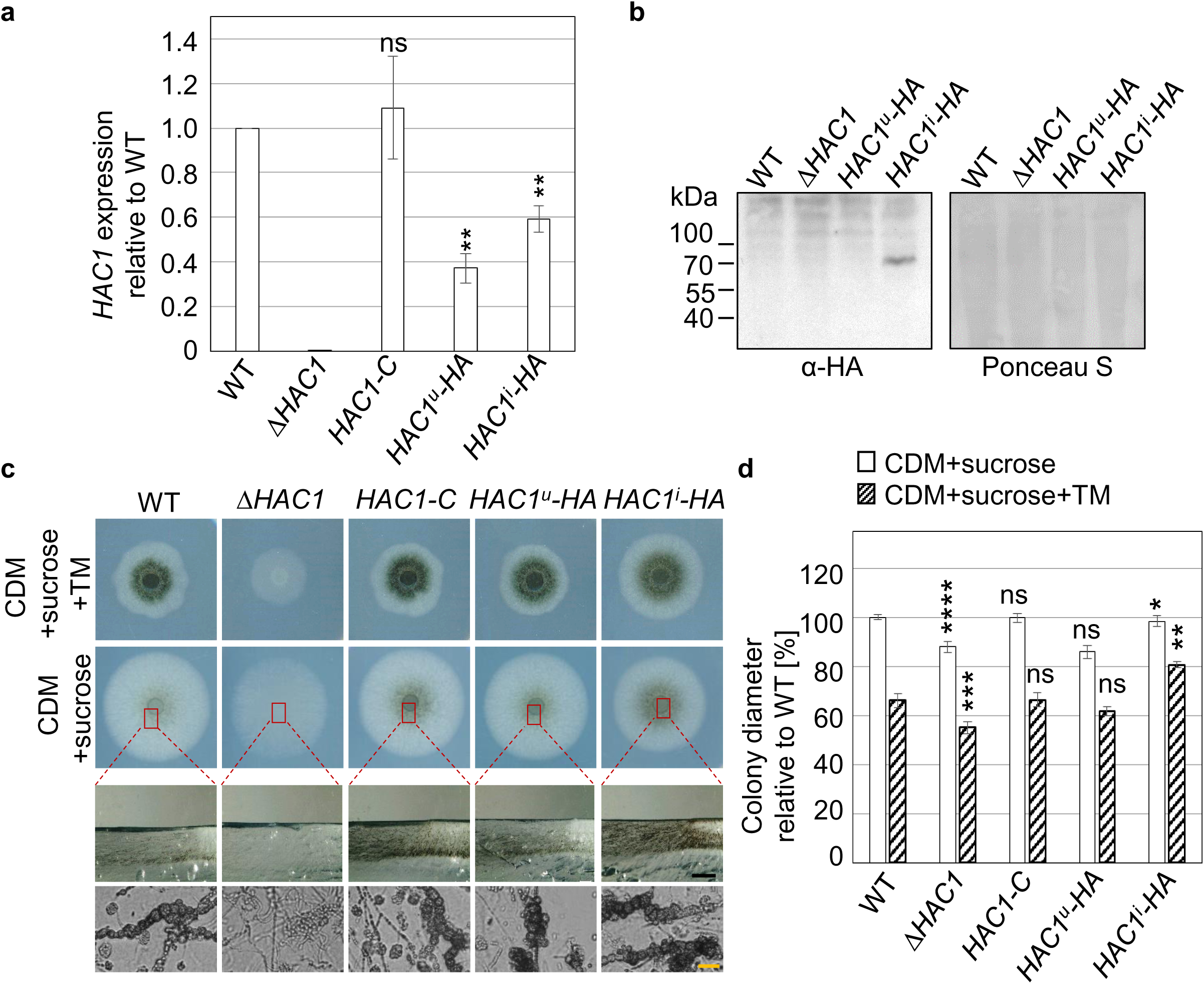
*V. dahliae HAC1* supports growth and is essential for microsclerotia formation. *Verticillium dahliae* wildtype (WT), *HAC1* deletion (Δ*HAC1*), and complementation (*HAC1-C*) strains, as well as strains expressing ectopically integrated *HAC1* mRNA splice variants fused to *HA* at the 3’-end in the Δ*HAC1* strain (*HAC1^u^-HA*; *HAC1^i^-HA*) were compared. (**a**) Quantification of *HAC1* gene expression by primers targeting both *HAC1* mRNA splice variants. Mean values of two independent experiments with standard deviations normalized to wildtype and reference genes *H2A* and *EIF2B* are shown (**p<0.01, ns = non-significant). *HAC1* gene expression is decreased in *HAC1^u^-HA* and *HAC1^i^-HA*. (**b**) Detection of Hac1 proteins in immunoblots using HA-specific antibodies. Ponceau S staining served as a loading control. A strong signal is only visible at ∼70 kDa for tagged Hac1 in the constitutively spliced *HAC1^i^-HA* strain instead of the predicted 46 kDA, whereas there was no specific band for the predicted Hac1^u^-HA protein (59 kDa). (**c**) Microsclerotia formation *ex planta.* Wildtype-like microsclerotia were formed by all strains except Δ*HAC1* 10 d after spot inoculation on CDM with sucrose with or without tunicamycin (TM = 1 µg/ml). Neither melanized nor unmelanized microsclerotia were observed for Δ*HAC1* in cross sections of the colony centres (red boxes/dashed lines) or microscopy of fungal material (bottom). Ectopically expressed *HAC1-C* and *HAC1^i^-HA* strains produce increased levels (Black scale bar = 1 mm, yellow scale bar = 20 µm). (**d**) Quantification of vegetative growth 10 d after spot inoculation. Δ*HAC1* displayed reduced growth that is restored to wildtype-levels in *HAC1-C* and *HAC1^u^-HA*, whereas *HAC1^i^-HA* displayed a decrease in growth under non-stress conditions and increased growth upon supplementation of tunicamycin (TM). Mean values of three independent experiments with standard deviations relative to wildtype are shown (*p<0.05; **p<0.01; ***p<0.001, ****p = 0, ns = non-significant, n≥2).

Alterations in growth and development of the constructed strains *ex planta* were compared to wildtype. Δ*HAC1* colonies produce less aerial mycelium and appear more transparent. Under non-stress conditions reduced growth was observed for Δ*HAC1* (15%) and *HAC1^i^-HA* (10%) in comparison to the wildtype strain (Fig. 2c, d). Under DTT-induced ER stress conditions growth of Δ*HAC1* was decreased (18% lower), whereas *HAC1^i^-HA* displayed relatively increased colony diameters (17% higher) compared to wildtype. The expression of the ectopically integrated *HAC1* gene in *HAC1-C* or the *HAC1* gene lacking the conventional intron in *HAC1^u^-HA* in the deletion background resulted in wildtype-like growth under stress- and non-stress conditions. This suggests, that presence of the unconventionally spliced *HAC1* mRNA in the *HAC1^i^-HA* strain enables a more efficient response to ER stress.

For Δ*HAC1* no melanization of the colony centres was observed during growth on any tested medium. Cross sections and microscopy of fungal material from Δ*HAC1* colonies grown on different media revealed the absence of microsclerotia, whereas ectopic integration resulted in increased resting structure occurrence for the *HAC1-C* and *HAC1^i^-HA* strains as exemplified for minimal medium with sucrose as carbon source (Fig. 2c). Wildtype-like microsclerotia frequencies were observed in the *HAC1^u^-HA* strain. The absence of microsclerotia in Δ*HAC1* and *vice versa* increased melanization in *HAC1^i^-HA* corroborate a regulatory function of *HAC1* in resting structure formation.

### Virulence of *V. dahliae* depends on the unfolded protein response transcription factor Hac1

As mentioned above, the unfolded protein response controls the capacity and quality of ER-mediated secretion. It is currently unknown whether the UPR regulator Hac1 is important for fungal pathogens that colonize plant roots. Therefore, we compared root colonization by a UPR defective *V. dahliae HAC1* deletion strain versus the wildtype strain. A *HAC1* deletion strain expressing ectopically integrated *GFP* under control of the *gpdA* promoter was constructed for monitoring colonization. Three-week-old *Arabidopsis thaliana* plants inoculated with the same number of spores obtained from Δ*HAC1 OE-GFP* and wildtype carrying the same construct (WT *OE-GFP^NAT^*) were compared. Fungal colonization behaviour was studied after five to seven days. Overall, less hyphae on the root surface were present for the Δ*HAC1 OE-GFP* strain compared to wildtype (Fig. 3a). The formation of swollen hyphae and a change in growth direction indicate penetration sites. These features were observed in the absence of *HAC1* similarly to wildtype, as was hyphal growth after invasion of the root outer layer. Therefore, *V. dahliae HAC1* is not required for penetration of the root and the mutant is not blocked directly after invasion of the root cortex, but *HAC1* supports the first contact with the host, which is the initial colonization of the root surface.

**Figure 3:**
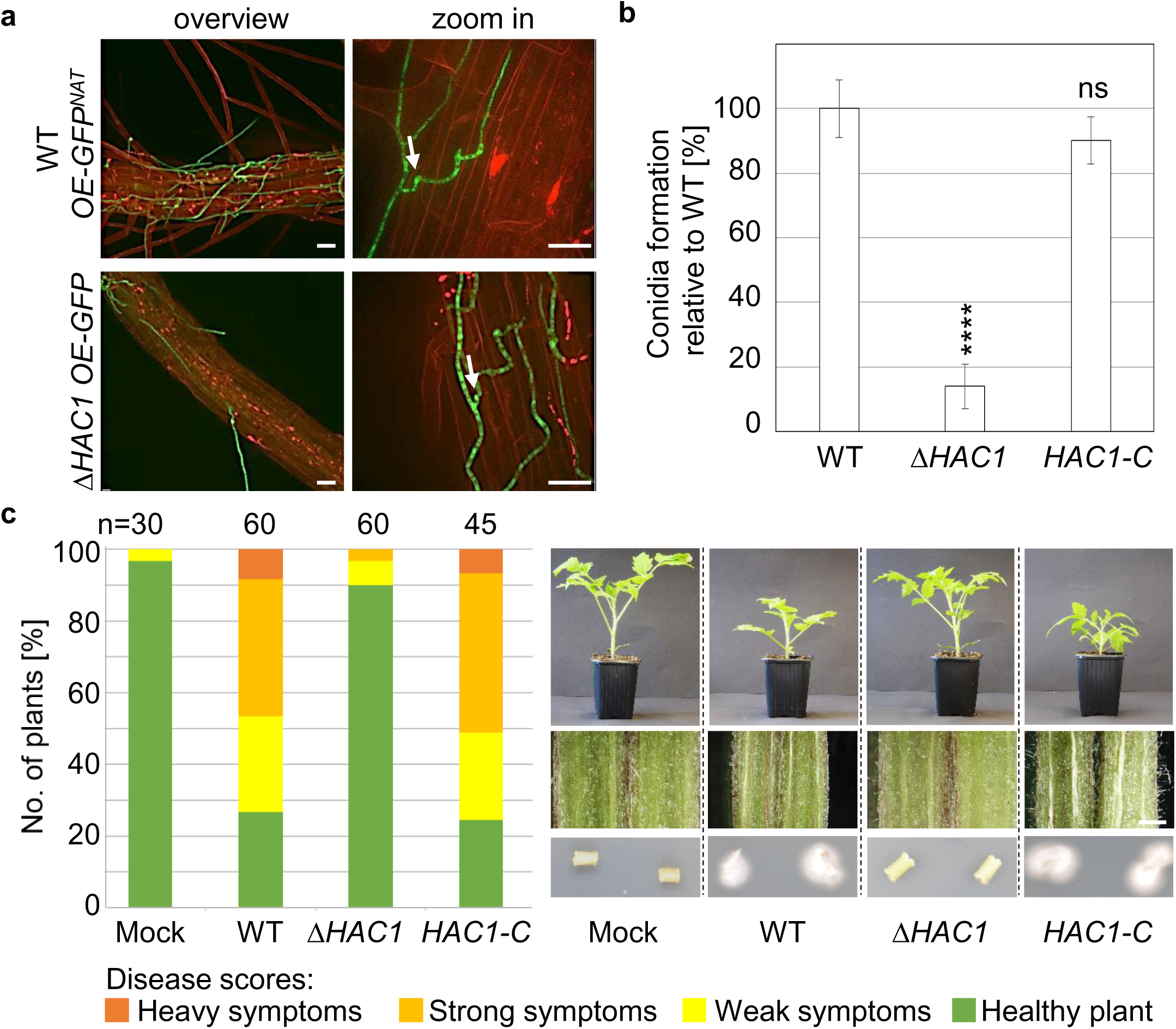
*V. dahliae HAC1* is required for initial plant root colonization, conidia formation, and induction of disease symptoms in tomato plants. *Verticillium dahliae* wildtype (WT), *HAC1* deletion (Δ*HAC1*), and complementation (*HAC1-C*) strains, as well as wildtype and *HAC1* deletion strains constitutively expressing ectopic GFP (WT *OE-GFP^NAT^*, Δ*HAC1 OE-GFP*) were compared. (**a**) *V. dahliae* colonization of *Arabidopsis thaliana* roots. Fluorescence confocal microscopy was performed 7 d post inoculation of roots with the same numbers of spores from wildtype WT *OE-GFP^NAT^* or Δ*HAC1 OE-GFP* with four plants per strain in two independent experiments. Δ*HAC1 OE-GFP* displays reduced propagation on the root surface, but was able to penetrate roots (white arrows) and colonize the root cortex (Scale bar = 20 µm). (**b**) Quantification of conidiation 5 d post inoculation of liquid simulated xylem medium revealed reduced Δ*HAC1* conidiation compared to wildtype. Mean values of four independent experiments with standard deviations relative to wildtype are shown (****p = 0, ns = non-significant, n≥4). (**c**) Pathogenicity test of Δ*HAC1* mutant compared to wildtype towards *Solanum lycopersicum*. Representative plants and hypocotyl dissections 21 d after root dipping into distilled water control (Mock), or same numbers of spores obtained from different strains are shown (Scale bar = 1 mm). Relative amount of plants with certain disease scores from two independent experiments are displayed in the stack diagram (n = total number of evaluated plants). Δ*HAC1* treated plants showed mock-like hypocotyl coloration, disease symptoms in only few plants and no fungal outgrowth from surface sterilized stem sections after 7 d (bottom).

*V. dahliae* forms conidiospores within the plant vascular system for spreading and systemic colonization. The *HAC1* deletion strain displayed significantly reduced conidiospore numbers with about 14% relative to wildtype after five days in liquid simulated xylem medium (Fig. 3b). Conidiospore levels were restored in the *HAC1-C* strain. This suggests a further role for *HAC1* function in subsequent steps of plant colonization, and this was investigated by tomato infection experiments. Δ*HAC1* induced significantly less severe disease symptoms than wildtype 21 days after inoculation with conidiospores, resulting in approximately 90% healthy plants (Fig. 3c). None of the Δ*HAC1* inoculated plants showed heavy symptoms, and no hypocotyl discolorations were observed for any plant. In addition, no fungal outgrowth was observed for Δ*HAC1* from stems of treated plants (Fig. 3c). The less virulent *in planta* phenotype of the Δ*HAC1* strain was complemented by *HAC1-C* regarding induction of overall disease symptoms, discoloration of the hypocotyl and fungal outgrowth from stem sections. These results support the conclusion that a functional UPR regulator Hac1 is required for the initial colonization of the host root surface and sporulation as prerequisite for fungal propagation within the plant. Hac1 is therefore required for induction of severe disease symptoms in tomato.

### *V. dahliae* development and plant disease symptom induction require pheromone response MAP kinase activities independently from the Ham5 scaffold

The unfolded protein response participates in cross regulation with the pheromone response MAPK pathway in the dimorphic plant pathogen *U. maydis* and this control prevents hypervirulence and maintains fungal biotrophy in the plant (Di Stasio *et al*., 2009; Heimel *et al*., 2013; Schmitz *et al*., 2019). Pheromone response MAPK pathways integrate the sensing of extracellular signals and activate downstream pathways that control pathogenic differentiation and fungal virulence for numerous fungi including *V. dahliae* (Jiang *et al*., 2018). Pathways are often assembled by scaffold proteins such as the yeast pheromone scaffold protein Ste5 to maintain specificity and prevent crosstalk between signalling pathways. The scaffold proteins Ham5/HamE of the pheromone response MAPK pathways of *Neurospora crassa* or *Aspergillus nidulans* are required for specific fungal developmental programs (Dettmann *et al*., 2014; Jonkers *et al*., 2014; Frawley *et al*., 2018). To date, corresponding scaffold proteins have not yet been analysed in plant pathogenic fungi and it is unknown whether they are required for pheromone response MAPK pathway mediated virulence.

*V. dahliae HAM5* was identified by reciprocal BLAST and the deduced protein shows 53% amino acid identity with the *N. crassa* scaffold homolog *HAM-5*. The 4906 bp *HAM5* pre-mRNA encodes a 1553 aa protein containing N-terminal WD40 repeats and a coiled-coil domain at the C-terminus. The corresponding proteins of related filamentous fungi are similar in length and show conserved protein domains. *V. dahliae MEK2* encodes a 522 aa MAP2K and *VMK1* encodes a 355 aa MAPK, and both include ATP binding and serine/threonine-protein kinase active sites (Fig. 4).

**Figure 4:**
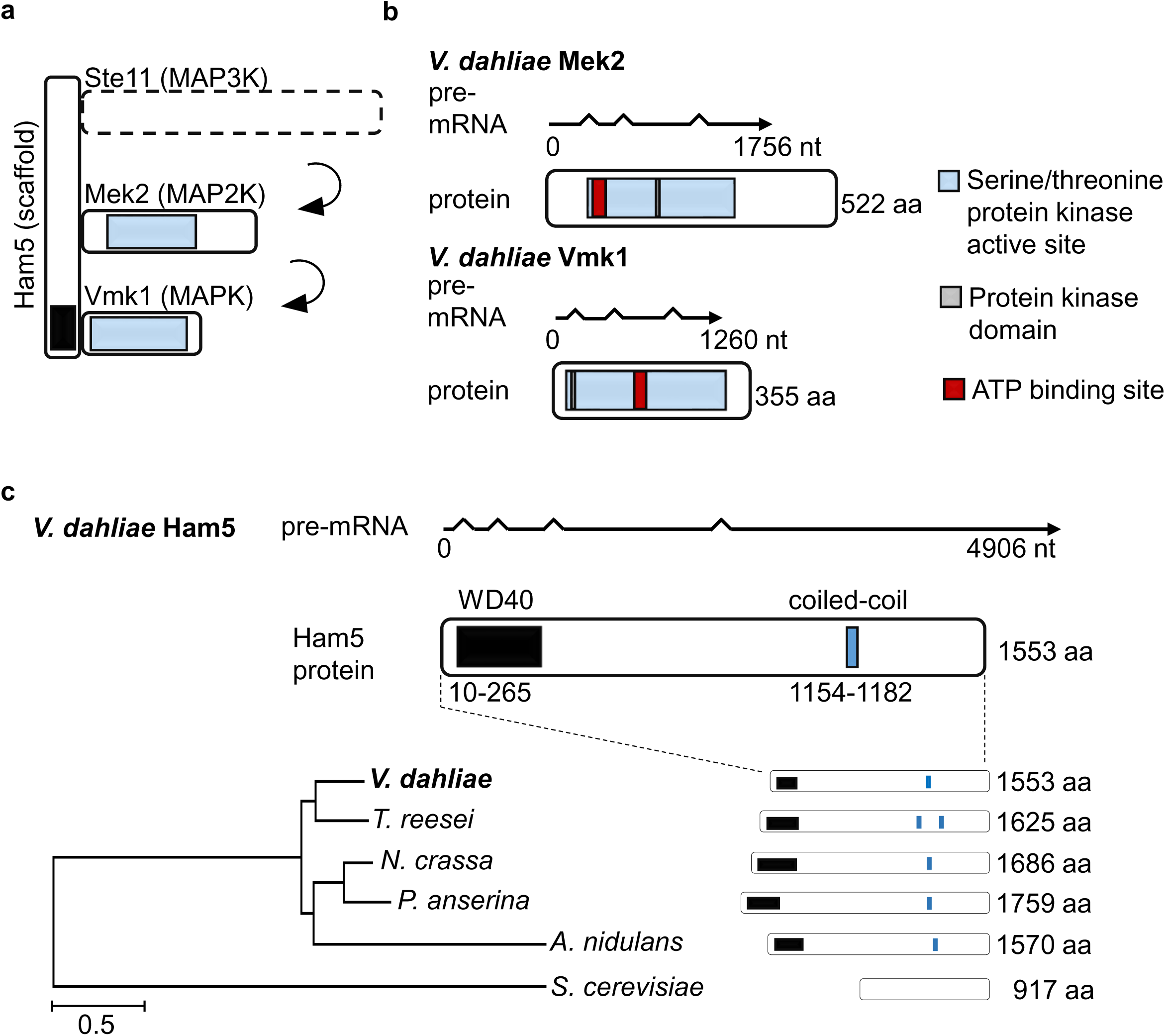
Pheromone response mitogen activated protein (MAP) kinase module of *V. dahliae.* Intron-exon boundaries and resulting open reading frames (ORF) were confirmed by PCR amplification and sequencing of wildtype cDNA. (**a**) Architecture of the *V. dahliae* MAPK module including Ham5 scaffold. (**b**) Transcript structures and deduced protein kinase domains of *V. dahliae* MAPK2 Mek2 (left) and MAPK Vmk1 (right). The Mek2 ORF results in 522 amino acids (aa) with protein kinase domain (blue bar: 67-332 aa; IPR000719) including active serine/threonine site residues (grey: 186-198aa, IPR008271) and ATP binding site (red: 73-96 aa; IPR017441). Vmk1 is smaller and consists of 355 aa (blue: protein kinase domain: 23-311 aa, IPR000719; grey: active site serine/threonine residues: 143-155 aa, IPR008271; red: ATP binding site: 29-53aa; IPR017441). (**c**) *HAM5* (*VDAG_JR2_Chr4g07170a*) transcripts and deduced protein domains. The 1553 aa Ham5 scaffold contains WD40 repeats at the N-terminus (black: 10-265 aa, IPR015943) and a coiled-coil domain (blue: 1154-1182 aa). Related Ham5-like proteins of other fungi are depicted in a phylogenetic tree (ClustalW algorithm) with *Trichoderma reesei* HAM-5 (AKN58846.1), *Neurospora crassa* HAM-5 (XP_011393509.1)*, Podospora anserina* IDC1 (ABJ96338.2), *Aspergillus nidulans* HamE (AN2701), and *Saccharomyces cerevisiae* Ste5 (NP_010388.1) (Scale bar = average number of amino acid substitutions per site).

The role of the scaffold Ham5 in the Vmk1 MAPK pathway for fungal development and virulence was compared to *MEK2* and *VMK1* through evaluation of the corresponding single and double deletion strains. The *HAM5* deletion strain revealed wildtype-like growth and resting structure formation *ex planta* under stress and non-stress conditions, exemplified for CDM with sucrose or cellulose 9 days after inoculation (Fig. 5a). *MEK2* and *VMK1* single and double deletion strains with *HAM5* exhibited a 10% decrease in growth 9 days after inoculation, which was restored in the complementation strains *MEK2-C* and *VMK1-C* (Fig. 5a, b). In addition, Δ*MEK2*, Δ*VMK1*, Δ*HAM5*Δ*MEK2,* and Δ*HAM5*Δ*VMK1* strains formed less microsclerotia on minimal medium with cellulose, and melanization of the colonies was reduced to less than half, whereas Δ*HAM5* and *HAM5-C* strains displayed wildtype-like melanization. Phenotypes were restored by ectopic integration of the corresponding wildtype gene in *MEK2-C* and *VMK1-C* strains (Fig. 5a, b).

**Figure 5:**
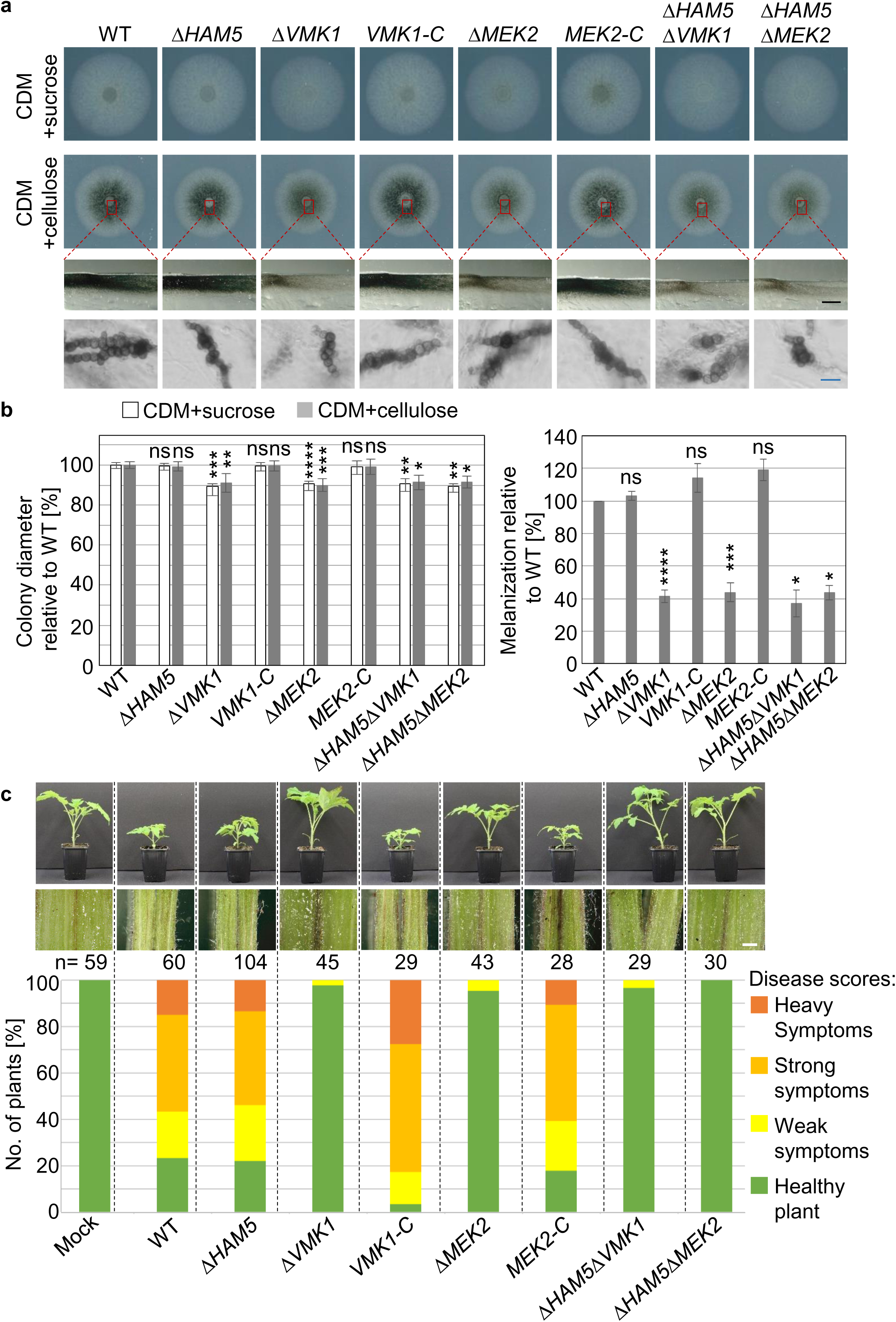
*V. dahliae* Vmk1 and Mek2 kinases but not the Ham5 scaffold are required for efficient melanization and virulence. The *HAM5* deletion strain (Δ*HAM5*) was compared to wildtype (WT), *VMK1* and *MEK2* single deletions (Δ*VMK1;* Δ*MEK2*) with respective complementation strains (*VMK1-C*; *MEK2-C*), as well as Δ*HAM5*Δ*VMK1* and Δ*HAM5*Δ*MEK2* double deletions. (**a**) Vegetative growth and melanization *ex planta.* 50 000 freshly harvested spores were spot inoculated on CDM with either sucrose or cellulose and incubated at 25°C for 9 d. Cross sections of colony centres (red boxes/dashed lines) and microscopy of fungal material scrapped from colony centres grown on CDM with cellulose are shown (black scale bar: 1 mm, blue scale bar: 20 µm). (**b**) **left:** Growth quantification 9 d after spot inoculation on CDM with sucrose or cellulose. Δ*HAM5* displays wildtype-like growth, whereas other single and double deletion strains are slightly repressed in growth. Mean values of three independent experiments with standard deviations relative to wildtype are shown (*p<0.05; **p<0.01; ***p<0.001, ****p = 0, n≥3). **right:** Melanization of colonies grown on CDM with cellulose relative to wildtype. Whereas Δ*HAM5* showed wildtype-like melanization, other single and double deletion strains displayed a decrease to about 40%. Mean values of two independent experiments with standard deviations relative to wildtype are shown (*p<0.05; ***p<0.001, ****p = 0, ns = non-significant, n≥2). (**c**) Pathogenicity test of MAPK pathway mutants towards *Solanum lycopersicum*. Representative plants and hypocotyl dissections 21 d after root dipping into distilled water as control (Mock), or the same numbers of spores obtained from different strains are shown (Scale bar = 1 mm). The relative number of plants with certain disease scores are displayed in the stack diagram (total number of evaluated plants = n). Δ*HAM5* induced wildtype-like disease symptoms, whereas Δ*MEK2*, Δ*VMK1*, Δ*HAM5*Δ*MEK2,* or Δ*HAM5*Δ*VMK1* treated plants were comparable to mock plants, and show no hypocotyl discolorations.

The impact of *V. dahliae* Ham5 was investigated in tomato plant infection experiments and compared to the core MAPK components Mek2 and Vmk1 (Fig. 5c). Symptoms of tomato plants inoculated with spores from Δ*MEK2*, Δ*VMK1*, Δ*HAM5*Δ*MEK2,* or Δ*HAM5*Δ*VMK1* strains were comparable to mock plants after 21 days, and no hypocotyl discolorations were observed (Fig. 5c). The avirulent *in planta* phenotypes were complemented in experiments with *VMK1-C and MEK2-C*. Disease symptoms induced by the Δ*HAM5* strain were indistinguishable from the wildtype control and plants displayed severe stunting and discoloration of the vascular tissue (Fig. 5c).

These data corroborate that the requirement of the MAPK cascade components Vmk1 and Mek2 for virulence, growth and microsclerotia formation is independent of the presence of Ham5 and its isolation function as scaffold protein in *V. dahliae*. These results highlight the potential for crosstalk between UPR and this MAPK module signalling pathway, given that both promote fungal virulence, growth and microsclerotia formation.

### The *V. dahliae* oleate Δ12-fatty acid desaturase Ode1 promotes fungal differentiation with only a minor impact on virulence

Activation of the yeast pheromone response MAPK pathway relies on the perception of peptide pheromones (Bardwell, 2005). To coordinate their development, several filamentous fungi produce oxylipins instead of pheromones as signalling molecules, which are as well connected to host-fungus communication (Calvo *et al*., 2001; Brodhagen *et al*., 2008; Brodhun *et al*., 2009; Reverberi *et al*., 2010; Scala *et al*., 2014; Fischer & Keller, 2016; Patkar & Naqvi, 2017). Secretion of these lipid metabolites potentially relies on the UPR control of the capacity and quality of ER mediated secretion. We examined functions of *V. dahliae* oxylipins as potential signals for development and virulence, and focused an oleate Δ12-fatty acid desaturase catalysing the oxidation of oleic acid to linoleic acid, the major precursor of fungal oxylipins. The impact of the oleate Δ12-fatty acid desaturase OdeA on differentiation was described in different Aspergilli, but has not yet been examined in plant pathogens (Calvo *et al*., 2001; Chang *et al*., 2004; Wilson *et al*., 2004).

*V. dahliae ODE1* was identified as the homolog to *A. nidulans odeA* by reciprocal BLAST search of the amino acid sequences with 66% aa sequence identity of the deduced proteins. The Ode1 protein with a length of 481 aa and a predicted molecular weight of 54 kDa contains two fatty acid desaturase (FAD) domains (Fig. 6a). The cytosolic catalytic centre of Δ12-fatty acid desaturases is formed by conserved N- and C-terminal histidine clusters described as FAD domains and iron atoms provided by the membrane-bound donor cytochrome b_5_ (Los & Murata, 1998). *V. dahliae* Ode1 was predicted as a transmembrane protein with four hydrophobic transmembrane helices, two short non-cytosolic regions, and three hydrophilic cytosolic regions, including the N- and C-termini of the protein (Fig. 6b).

**Figure 6:**
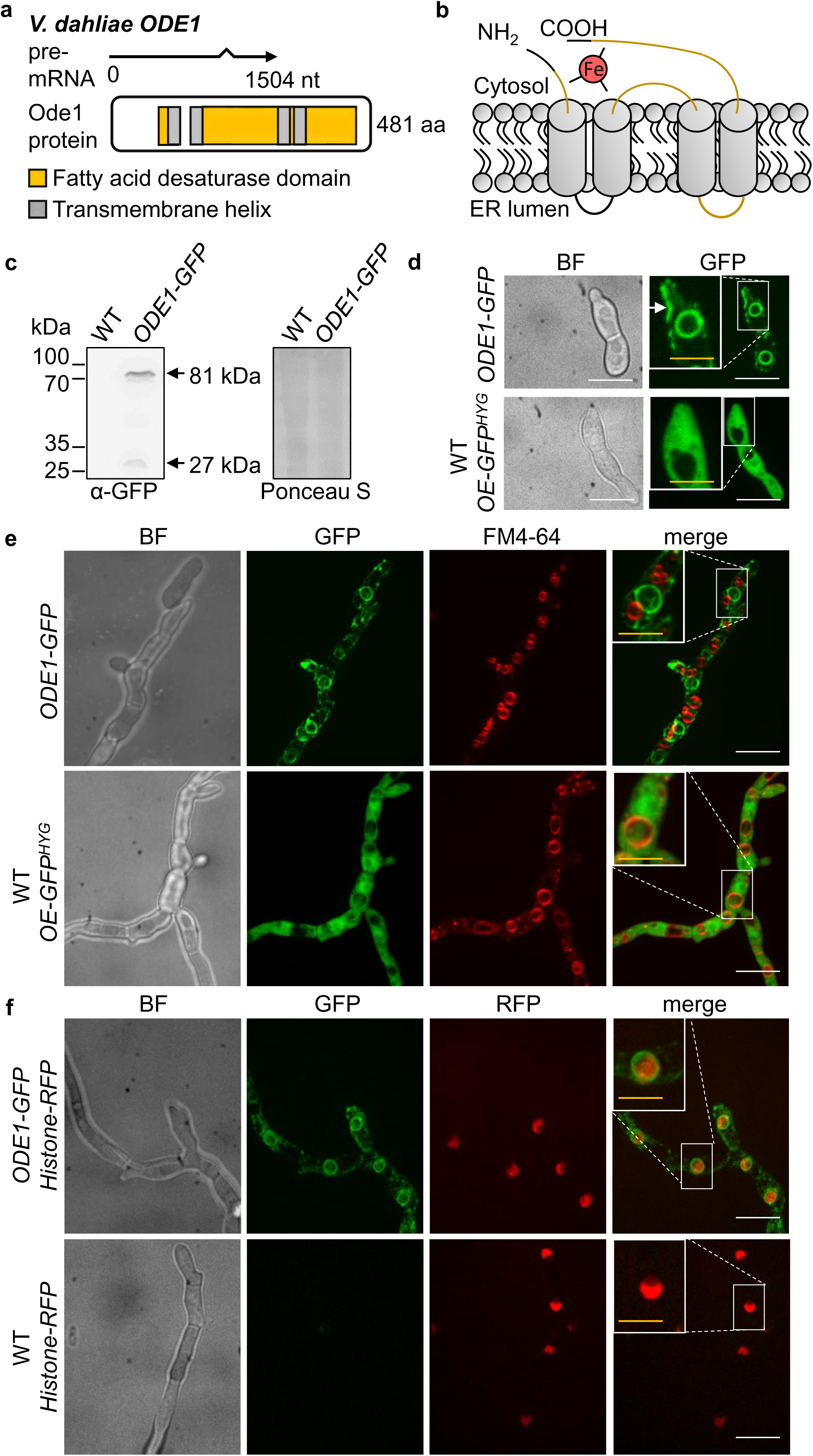
*V. dahliae* Ode1 is localized to plasma and nuclear membranes. The *ODE1-GFP* strain harbours a functional fusion at the endogenous locus under control of the native promoter and terminator. This strain with or without RFP tagged histone H2B, and wildtype constitutively expressing free GFP (WT *OE-GFP^HYG^*) or RFP tagged histone H2B (JR2 *Histone-RFP*) were compared. (**a**) *Verticillium dahliae* oleate Δ12-fatty acid desaturase encoding *ODE1* intron-exon boundaries (1504 bp) were confirmed by PCR amplification from wildtype cDNA and sequencing. The deduced Ode1 protein (481 aa) contains two fatty acid desaturase domains (FAD; yellow; 77-112 aa, IPR021863; 138-424 aa; IPR005804), and four putative transmembrane helices (grey; 105-124 aa, 136-157 aa, 300-319 aa, 331-350 aa; Phobius). (**b**) Scheme of the predicted Ode1 protein structure. The N- and C-termini of the transmembrane protein are directed to the cytosol. An iron atom (Fe) and the FAD domains (yellow) form the catalytic centre. (**c**) Detection of Ode1-GFP (81 kDa) in immunoblot analysis using GFP-specific antibody. Ponceau S staining served as loading control. (**d**) Fluorescence microscopy revealed Ode1-GFP subcellular localization to membranes of round cell organelles and close to hyphal tips (white arrow) of growing hyphae 12 h after inoculation (Scale bar = 10 µm). (**e**) Ode1-GFP is not localized to FM4-64 red stained vacuole membranes 16 h post inoculation (scale bar = 10 µm). (**f**) Ode1-GFP is localized to membranes surrounding red nuclei in the *ODE1-GFP Histone-RFP* strain. Fluorescence microscopy 16 h post inoculation (yellow scale bar = 20 µm, white scale bar = 10 µm).

*V. dahliae ODE1* deletion and complementation strains harbouring *ODE1* C-terminally fused to GFP at the endogenous locus under control of the native promoter and terminator were constructed. The predicted molecular weight for Ode1 fused to GFP with 81 kDa was confirmed by immunoblots (Fig. 6c). Fluorescence microscopy of young hyphae revealed that the Ode1-GFP protein is primarily localized to ER membranes surrounding the nucleus and to a minor extend to plasma membranes and close to tips of growing hyphae resembling perinuclear and cortical ER structures (Fig. 6d-f).

The impact of the *V. dahliae* oleate Δ12-fatty acid desaturase Ode1 on fungal growth and differentiation was examined by comparing the *ODE1* deletion to the wildtype strain under different physiological and membrane stress inducing conditions. Vegetative growth of the Δ*ODE1* strain was significantly decreased with or without stressors (Fig. 7a). The *ODE1* deletion strain displayed the most severe decrease in vegetative growth on medium containing cellulose, with a ∼50% reduction of the colony diameter after 9 days. This defect was complemented by *ODE1-GFP* (Fig. 7a, b). The observed growth defect of Δ*ODE1* was partially compensated by supplementation of media with linoleic acid, resulting in a relative colony diameter of about 70% (Fig. 7b). Reduced melanization correlated with the decrease in growth of colonies formed by Δ*ODE1* on cellulose medium. Colony cross sections and microscopy of fungal material from colony centres revealed the formation of wildtype-like microsclerotia with regard to size, shape and melanization by the Δ*ODE1* strain (Fig. 7a).

**Figure 7:**
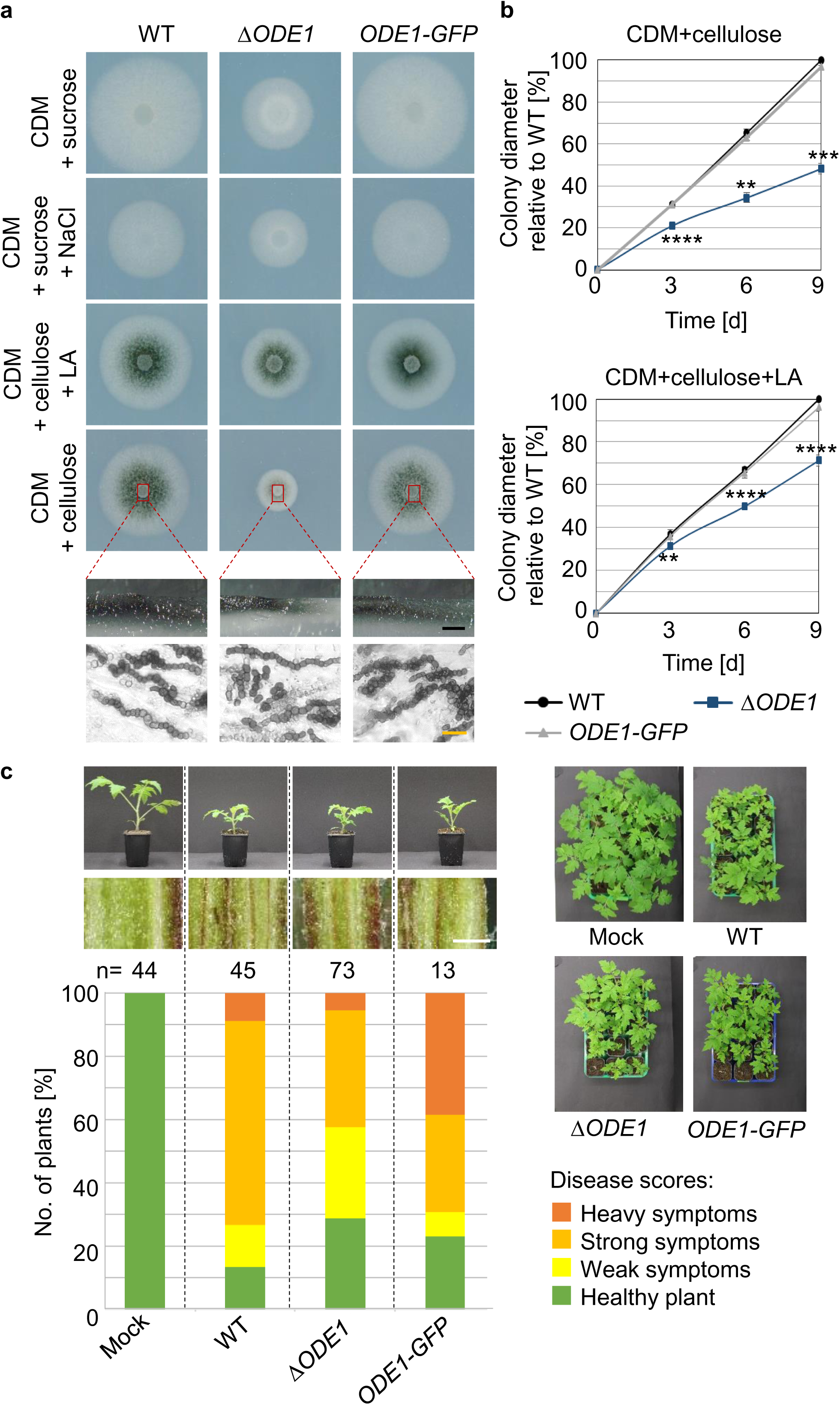
*V. dahliae ODE1* contributes primarily to vegetative growth with only minor effects on plant disease symptoms. The *Verticillium dahliae ODE1* deletion strain (Δ*ODE1*) was compared to wildtype (WT) and the complementation strain harbouring *ODE1-GFP* at the endogenous locus under control of the native promoter and terminator. (**a**) Vegetative growth and microsclerotia formation *ex planta* 9 d after spot inoculation on CDM supplemented with NaCl (0.5 M), CDM with either sucrose or cellulose, and CDM with cellulose supplemented with linoleic acid (LA, 0.125 mg/ml). Δ*ODE1* formed smaller colonies on all tested media. Cross sections of colonies grown on CDM with cellulose (red boxes/ dashed lines) and microscopy of fungal material (bottom) showed wildtype-like microsclerotia formation of Δ*ODE1* (black scale bar = 1 mm, yellow scale bar = 20 µm). (**b**) Growth quantification 3, 6, and 9 d after spot inoculation on CDM with cellulose with or without linoleic acid (LA, 0.125 mg/ml). Δ*ODE1* displayed about 50% decreased growth 9 d after spot inoculation on CDM with cellulose. LA supplementation partially complements the growth defect. Mean values and standard deviations relative to wildtype of two independent experiments are shown (**p<0.01; ***p<0.001, ****p = 0, n≥3) (**c**) Pathogenicity test of *ODE1* mutants towards *Solanum lycopersicum*. Representative plants and hypocotyl dissections 21 d after root dipping into distilled water control (Mock), or same number of spores obtained from different strains are shown (Scale bar = 1 mm). Relative amount of plants with certain disease scores from three independent experiments are displayed in the stack diagram (n = total number of evaluated plants). Δ*ODE1* infection resulted in only a minor decrease in disease symptom induction.

Tomato plant infection experiments addressed the impact of *ODE1* on the virulence of *V. dahliae*. Tomato plants inoculated with Δ*ODE1* spores showed only slightly reduced numbers of plants with disease symptoms after 21 days compared to stunting and hypocotyl discolorations of wildtype infected plants (Fig. 7c). This suggests a strong contribution of the single gene *ODE1* to fungal growth but only a minor contribution to the virulence of *V. dahliae*.

These data hint at a complex interplay between biosynthesis of linoleic acids as precursors of oxylipins, the secretion capacity moderating UPR, and the scaffold-independent pheromone response MAPK pathway, which perceives external signals and activates downstream pathways. This interplay results in promotion or reduction of fungal growth and influences the outcome of the interaction of *V. dahliae* with its host plant.

## Discussion

The unfolded protein response and the Ham5 scaffold-independent pheromone response MAPK pathway form important distinct signalling hubs for *V. dahliae* plant pathogenicity. The UPR regulator Hac1 is essential for induction of resting structure formation and conidiation, the first contact with the plant roots and propagation within the plant. The pheromone response MAPK pathway does not require the scaffold Ham5 to support maturation and melanization of microsclerotia as well as prompting disease symptoms as severe stunting and discoloration of the vascular tissue. The linoleic acid synthesizing oleate Δ12-fatty acid desaturase Ode1 primarily affects fungal growth with only a small contribution to virulence of *V. dahliae*. The interplay between these signalling pathway components in *V. dahliae* growth, development and induction of plant disease is summarized in Fig. 8.

**Figure 8:**
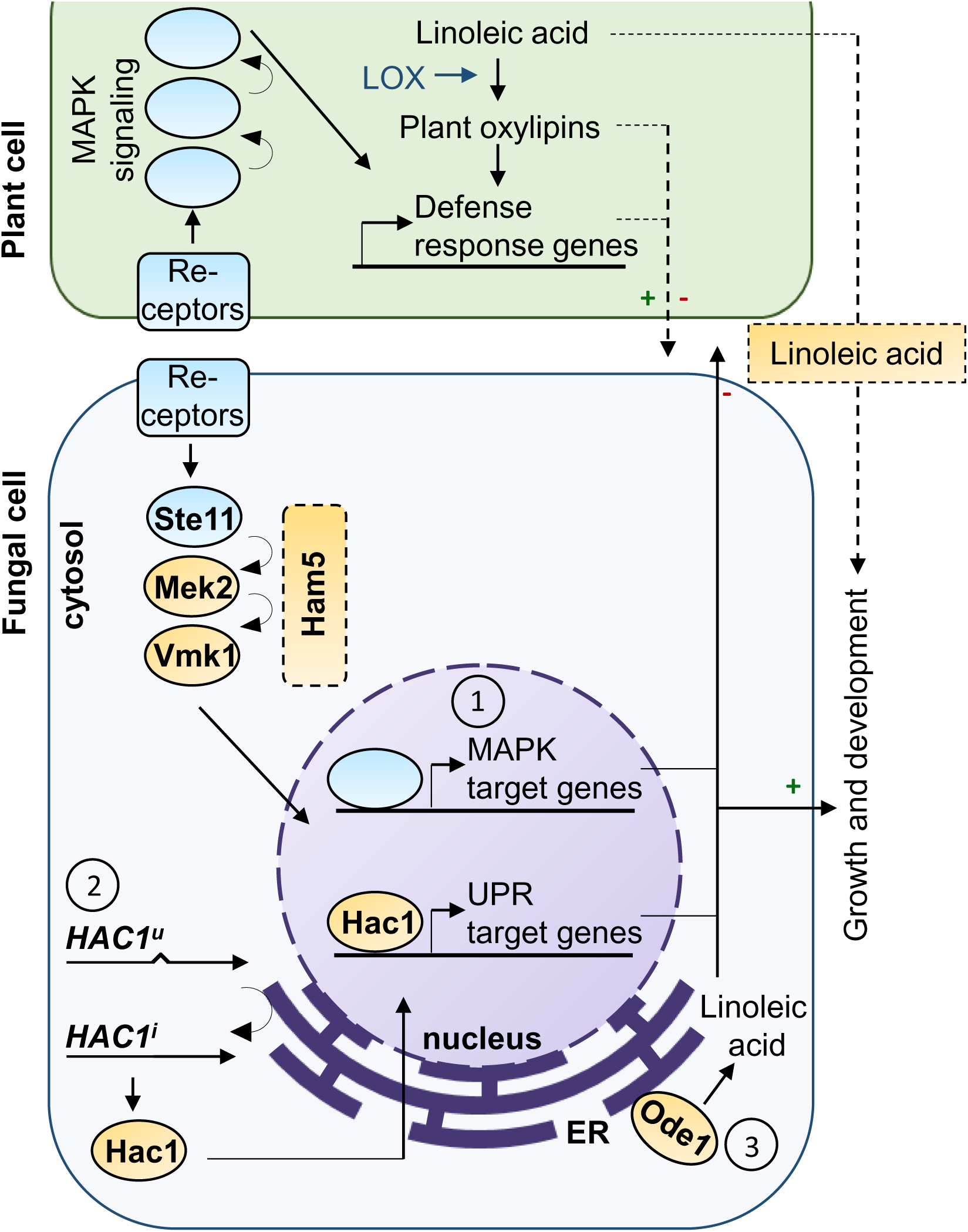
Model of *V. dahliae* signalling pathways and their functions in development and induction of disease symptoms. ① The unspliced mRNA of the UPR regulator Hac1 (*HAC1^u^*) is unconventionally spliced (arrow close to ER membrane), and the *HAC1^i^* mRNA is translated into the bZIP transcription factor Hac1, which regulates UPR target genes. *V. dahliae HAC1* is involved in the ER stress response and vegetative growth under non-stress conditions, has a strong impact on conidiation, and is essential for the formation of microsclerotia (arrow/ green plus to “Growth and development”). *HAC1* is required for the virulence of *V. dahliae* (arrow/ red minus from fungal cell) and potentially regulates expression or secretion of effector proteins, which enables the fungus to circumvent plant defence responses (arrow with dashed line/ green plus from plant cell). ② The *V. dahliae* Vmk1 MAPK pathway components Mek2 (MAP2K) and Vmk1 (MAPK) positively control vegetative growth and microsclerotia formation (arrow/ green plus to “Growth and development”) and are required for induction of disease symptoms in plants (arrow/ red minus from fungal cell) independent from the scaffold homolog Ham5 in *V. dahliae.* ③ The oleate Δ12-fatty acid desaturase Ode1, which catalyses the synthesis of linoleic acid, supports fungal growth (arrow/ green plus to “Growth and development”). Ode1 deficiency has only minor effect on development of disease symptoms *in planta*, which might be due to the availability of plant linoleic acid to potentially to support fungal growth and differentiation (arrow with dashed line/ green plus from plant cell).

The *V. dahliae* UPR regulator Hac1 has conserved, as well as species-specific impacts on fungal differentiation and is important for virulence. The *V. dahliae HAC1* mRNA contains an unconventional intron with sequence and structural similarity to Ire1 spliced introns in other organisms. In contrast to wildtype-like *HAC1* gene expression levels in the complementation strain harbouring the entire *HAC1* gene, *HAC1^i^-HA* and *HAC1^u^-HA* strains harbouring either one of two *HAC1* splice variants without conventional intron in its genome displayed decreased expression. This might result from intron-mediated enhanced transcription of *HAC1*.

Only the Hac1 protein resulting from translation of unconventionally spliced mRNA, but not from the uninduced variant, produced sufficient protein amounts for detection by immunoblots. Translation of *HAC1^u^* mRNA might be blocked or result in an unstable protein due to similar mechanisms as described for other ascomycetes. In *S. cerevisiae* base-pairing interaction between unconventional intron and 5’UTR leads to inhibition of ribosomal translation (Chapman & Walter, 1997; Rüegsegger *et al*., 2001). Additionally, accelerated degradation of Hac1^u^ proteins was described in yeast (Di Santo *et al*., 2016). Due to shortened intron length in *HAC1^u^* mRNA of filamentous ascomycetes, a similar mechanism for translation inhibition as in yeast is not possible (Mulder & Nikolaev, 2009). Rather, an impact of upstream ORFs on translational repression of unspliced *HAC1* mRNA was assumed (Saloheimo *et al*., 2003; Mulder *et al*., 2004), but studies in *Aspergillus niger* revealed base-pairing of the 5’UTR of *HAC1* mRNA with an inverted repeat sequence as translation attenuation mechanism (Mulder & Nikolaev, 2009). Truncation of the 5’UTR was described to foster translation of *HAC1* mRNA upon ER stress in different Aspergilli, *T. reesei* and *A. brassisicola* (Saloheimo *et al*., 2003; Mulder *et al*., 2004; Joubert *et al*., 2011).

*V. dahliae* is viable in the presence of constitutively induced *HAC1* mRNA and does not require the presence of the uninduced variant. In contrast, regulatory roles of the unspliced mRNA homolog and the deduced protein were proposed for *U. maydis* (Heimel *et al*., 2013). Here, overexpression of the induced mRNA resulted in UPR hyperactivation and lethality unless unspliced mRNA was present in the cell (Heimel *et al*., 2013).

In *S. cerevisiae*, *U. maydis* and *C. neoformans* the UPR is not required for vegetative growth and sporulation in the absence of ER stress (Nikawa *et al*., 1996; Kaufman, 1999; Cheon *et al*., 2011; Heimel *et al*., 2013). In contrast, deletion of *HAC1* or genomic integration of the unconventionally spliced *HAC1* mRNA variant affects growth and conidiation without stress in different filamentous ascomycetes including plant pathogens (Mulder & Nikolaev, 2009; Joubert *et al*., 2011; Carvalho *et al*., 2012; Tang *et al*., 2015). A basal UPR activity under non-stress conditions has been observed in several fungi (Heimel, 2015). Constitutive activation of the UPR might alter the control of genes involved in growth and developmental processes (Heimel, 2015). The *V. dahliae HAC1* deletion strain was also generally impaired in growth and was not additionally impaired in response to tunicamycin. Expression of the induced *HAC1* splice variant improved the ability to cope with ER stress, although expression levels of the induced splice variant were lower compared to wildtype. Under non-stress conditions, *HAC1^i^-HA* displayed reduced growth, which might be explained by differential regulation of genes supporting growth in correlation to a hyperactive UPR. This idea is supported by the finding, that the induced mRNA variant of the UPR regulator could be amplified from cultures grown under non-stress conditions in *V. dahliae*.

*HAC1* of the vascular pathogen *V. dahliae* is not only required for growth and conidiation in the absence of typical ER stress inducing conditions, but is also essential for the formation of microsclerotia. *HAC1* deletion strains do not form these resting structures under any tested condition. Decreased expression of the ectopically integrated *HAC1* gene lacking the conventional intron in the deletion background was sufficient to complement the microsclerotia phenotype in *HAC1^u^-HA*. For *HAC1^i^-HA* increased microsclerotia formation was observed. One possible explanation for *HAC1* being essential for microsclerotia production is that the UPR is a checkpoint to induce resting structure formation after sensing of unfavourable conditions. The mechanisms activating this process in *V. dahliae* are not yet understood, even if several candidates were shown to influence microsclerotia production such as the pheromone response MAPK pathway components Vmk1 and Mek2. An essential protein for microsclerotia formation is the transcription factor Som1 (Bui *et al*., 2019). Som1 is involved in regulation of a subset of genes, such as the Verticillium transcription activators of adhesion Vta2 and Vta3, which are involved in adhesion and microsclerotia formation (Tran *et al*., 2014; Bui *et al*., 2019). In *S. cerevisiae* Hac1 is involved in regulation of flocculin genes and interacts with the general control of amino acid biosynthesis (Herzog *et al*., 2013), and ER stressed cells display increased flocculation (Scrimale *et al*., 2009). The absence of microsclerotia in Som1 and Hac1-deficient *V. dahliae* strains hints to a crosstalk between Som1 and the UPR during regulation of microsclerotia formation.

In *U. maydis* the Hac1 orthologue is required to induce biotrophic growth *in planta* (Heimel *et al*., 2010). In contrast, UPR components are involved in initial penetration of the plant surface by *M. oryzae* (Yi *et al*., 2009; Tang *et al*., 2015; Jiang *et al*., 2018). The *V. dahliae* UPR regulator Hac1 has a major impact on the fungal ability to colonize host plants, and *HAC1* is required for efficient colonization of the root surface as first step. The subsequent penetration and initial root cortex invasion does not require Hac1, which is reminiscent of the finding that initial penetration of the plant surface was unaffected in the appressorium-forming fungus *A. brassicicola* as well as in *U. maydis* (Heimel *et al*., 2010; Joubert *et al*., 2011).

Tomato plants treated with the *V. dahliae HAC1* deletion strain displayed severely decreased disease symptoms and fungal re-isolation from treated stems was not possible. Successful propagation within the host requires fungal spreading within the vascular system by conidiation, which is defective in the *HAC1* deletion strain. In addition, the UPR considerably contributes to the necessary adaptation of secretory capacities during host colonization of pathogens and the processing and secretion of fungal effectors in *U. maydis* (Richie *et al*., 2011; Heimel *et al*., 2013; Hampel *et al*., 2016; Pinter *et al*., 2019).

Similar to the UPR, the pheromone response MAPK pathway is required for virulence of *V. dahliae* and contributes to growth and microsclerotia formation. Different MAPK pathways can share components, like MAP kinases, adaptor proteins or upstream kinases. For certain cascades scaffold proteins are necessary to bring components in proximity and maintain pathway specificity by isolation (Schaeffer & Weber, 1999; Patterson *et al*., 2010). Scaffold proteins are described for the pheromone response MAPK pathway in different filamentous ascomycetes and support fusion of fungal cells (Dettmann *et al*., 2014; Jonkers *et al*., 2014; Frawley *et al*., 2018). The *N. crassa* scaffold protein HAM-5 assembles the MAPK cascade during chemotropic growth and positively influences growth and differentiation (Li *et al*., 2005; Aldabbous *et al*., 2010; Dettmann *et al*., 2014; Jonkers *et al*., 2014). The homologous *A. nidulans* scaffold protein HamE is required for sexual and asexual development and secondary metabolite production (Frawley *et al*., 2018). The role of homologous scaffold proteins was yet unstudied in plant pathogens. Our findings demonstrate that pheromone response MAPK cascade mediated growth, differentiation and virulence are independent from isolation of this pathway by the Ham5 scaffold protein in *V. dahliae*.

In the basidiomycete *U. maydis* the phosphatase Rok1 (regulator of Kpp2) is responsible for dephosphorylation of the partially redundant MAP kinases Kpp2 and Kpp6 (kinase PCR-product 2/6) (Di Stasio *et al*., 2009). *U. maydis rok1* deficient mutants displayed a hypervirulent phenotype on maize plants. A scaffold protein involved in isolation of the MAPK cascade was not described in *U. maydis*. Recently it was shown, that Rok1 activity is regulated and induced by the UPR pathway (Schmitz *et al*., 2019). The orthologous phosphatase in *V. dahliae* (*VDAG_JR2_Chr7g08960a*) with 803 aa harbours a dual specificity phosphatase domain (337-528 aa, IPR020422) and shows high similarity to corresponding proteins of filamentous ascomycetes, but is more distantly related to *U. maydis* Rok1 or *S. cerevisiae* Msg5 (Supplementary Figure S4). As in the basidiomycete *U. maydis* the corresponding protein of the filamentous ascomycete *M. oryzae* functions in dephosphorylation of the pheromone responsive MAPK (Wang *et al*., 2017). This observation suggests that future studies should investigate the possibility of a connection between UPR and pheromone response MAPK in the vascular ascomycete pathogen *V. dahliae*.

Biosynthesis and secretion of lipid metabolites involved in growth, differentiation and virulence of pathogenic fungi might be controlled by the pheromone response MAPK cascade and the ER homeostasis moderating UPR. In several cases, the virulence of plant pathogenic fungi depends on oxylipin signalling (Brodhun & Feussner, 2011). Characterization of the *V. dahliae* oleate Δ12-fatty acid desaturase Ode1, catalysing the biosynthesis of linoleic acid, revealed an important contribution to fungal growth with only minor impact on induction of disease symptoms. Similarly, a general decrease in growth was observed for *A. nidulans* and *Aspergillus parasiticus* upon deletion of homologous oleate Δ12-fatty acid desaturases (Calvo *et al*., 2001; Chang *et al*., 2004; Wilson *et al*., 2004). *V. dahliae ODE1* is dispensable for formation of wildtype-like resting structures, whereas deletion of the *A. parasiticus ODE1* homolog resulted in loss of sclerotia development (Chang *et al*., 2004; Wilson *et al*., 2004).

Despite its impact on growth, *V. dahliae* Ode1 is not required for induction of severe disease symptoms in tomato plants. It is possible, that different oleate Δ12-fatty acid desaturases encoded in the *V. dahliae* genome are responsible for linoleic acid biosynthesis during plant colonization and compensate for loss of Ode1 function. In addition, the *ODE1* deletion strain might be able to use plant-derived unsaturated fatty acids to compensate for the growth defect. Plant linoleic acid and the derived oxylipins are recognized as mimics of fungal signalling molecules and promote sporulation and mycotoxin production in Aspergilli (Burow *et al*., 1997; Calvo *et al*., 1999; Wilson *et al*., 2004; Brodhagen *et al*., 2008; Gao & Kolomiets, 2009; Horowitz Brown *et al*., 2009; Reverberi *et al*., 2010).

In conclusion, *V. dahliae* Hac1 regulated unfolded protein response and a scaffold-independent pheromone response MAPK pathway are important for both resting structure formation and plant infection. They are potentially interconnected and represent interesting targets to control growth, survival of microsclerotia in the soil and propagation in the plant as a means to reduce the impact on crops of this important vascular pathogen.

## Acknowledgments

The authors thank N. Scheiter for technical assistance, K. Heimel for fruitful discussions and carefully reading the manuscript, S. Balnojan for construction of the plasmid pME4815, J. Teer for construction of the plasmid pME4976, A. Nagel for construction of the VGB493/VGB494, A. Nagel, J. Teer and E. F. Hettwer for support as student research assistants, as well as M. Leonard and A. Höfer for support. This work was funded by the Deutsche Forschungsgemeinschaft (DFG) IRTG 2172 (JS, IM), an NSERC CREATE award (JWK) and DFG BR1502/15-1 (RH, GHB).

## Author contributions

Conceived and designed the experiment JS, RH, IM, JWK and GHB. Performed the experiments JS, RH, IM, and RB. Analysed the data JS, RH, GHB. Wrote the paper JS, RH, and GHB.

## Supporting Information

Additional information may be found in the online version of this article.

## Supporting Information Figures

**Figure S1** Southern hybridization of *V. dahliae HAM5, VMK1* and *MEK2* single and double deletion, and complementation strains.

**Figure S2** Southern hybridization of *V. dahliae HAC1* deletion, *HAC1* deletion with ectopic *GFP* overexpression, *HAC1-C* complementation, as well as *HAC1^u^-HA* and *HAC1^u^-HA* strains.

**Figure S3** Southern hybridization of *V. dahliae ODE1* deletion and *ODE1-GFP* complementation strains.

**Figure S4** Phylogeny of Rok1-like phosphatases.

## Supporting Information Tables

**Table S1** Verticillium strains constructed and used in this study.

**Table S2** Primers used in this study.

**Table S3** Plasmids constructed and used in this study.

**Table S4** The cDNA sequence of *V. dahliae* JR2 *HAC1^i^.*

**Table S5** The amino acid sequence of *V. dahliae* JR2 Hac1.

## Supporting Information Methods

Methods S1

