## Supporting Information for "Unfolded protein response and scaffold independent pheromone MAP kinase signalling control *Verticillium dahliae* growth, development and plant pathogenesis"

**Figure S1: Southern hybridization of *V. dahliae HAM5, VMK1* and *MEK2* single and double deletion, and complementation strains.** *V. dahliae* wildtype (WT), *HAM5, VMK1* and *MEK2* single (Δ*HAM5* transformant number 1 and 2; Δ*VMK1* transformant number 2 and 4*;* Δ*MEK2* transformant number 21 and 22) and double deletion (Δ*HAM5*Δ*VMK1;* Δ*HAM5*Δ*MEK2*), and complementation (*HAM5-C; VMK1-C; MEK2-C*) strains were tested. *HYG*$^{R}$*:* hygromycin resistance cassette under control of the *gpdA* promoter and a *trpC* terminator; *NAT*$^{R}$: nourseothricin resistance cassette under control of the *gpdA* promoter and a *trpC* terminator. Ectopic integration sites of complementation strains are indicated by // at the end of the endogenous locus and by a blue line for the ectopic locus. 3´flanking regions were used as probes for Southern hybridization (indicated in red). (**a**) Southern hybridization of *HAM5* single and double deletion and complementation strains with *Sac*I. **left:** Scheme of restriction sites (arrows) and expected fragment length. **right:** Predicted bands of 4.9 kb for wildtype, 8.1 kb for the deletion, and 2.2 kb and 8.1 kb for the complementation were observed. (**b**) Southern hybridization of *VMK1* single and double deletion and complementation strains with *Bgl*I. **left:** Scheme of restriction sites (arrows) and expected fragment length. **right:** Predicted bands of 2.6 kb for wildtype, 2.4 kb for the deletion, and 2.1 kb and 2.4 kb for the complementation were observed. (**c**) Southern hybridization of *MEK2* single and double deletion and complementation strains with *Nru*I. **left:** Scheme of restriction sites (arrows) and expected fragment length. **right:** Predicted bands of 5.2 kb for WT, 7.1 kb for the deletion, and 5.3 kb and 7.1 kb for the complementation were observed.

**Figure S2: Southern hybridization of *V. dahliae HAC1* deletion, *HAC1* deletion with ectopic *GFP* overexpression, *HAC1-C* complementation, as well as *HAC1*$^{u}$*-HA* and *HAC1*$^{i}$*-HA* strains.** *V. dahliae* wildtype (WT), *HAC1* deletion (Δ*HAC1* transformant number 1 and 23), complementation (*HAC1-C*) strains and strains harbouring either one of the two ectopically integrated *HAC1* mRNA splice variants (*HAC1*$^{u}$*-HA* transformant number 2 and 5; *HAC1*$^{i}$*-HA* transformant number 6 and 10), or the ectopically integrated *GFP* overexpression construct (Δ*HAC1 OE-GFP*) were tested. Genomic DNA was cut with *Sal*I restriction enzyme and the 3´ flanking region of *HAC1* was used as a probe. *Hyg*$^{R}$: hygromycin resistance cassette under control of the *gpdA* promoter and a *trpC* terminator; *Nat*$^{R}$: nourseothricin resistance cassette under control of the *gpdA* promoter and a *trpC* terminator. **left:** Scheme of restriction sites (arrows) and expected fragment length. Ectopic integration sites of complementation strains are indicated by // at the end of the endogenous locus and by a blue line for the ectopic locus. **right:** Predicted bands of 2.1 kb for wildtype, 3.2 kb for Δ*HAC1* and Δ*HAC1 OE-GFP,* and 3.2 kb and 1.8 kb for the *HAC1-C,* $HAC1^{u}$*-HA,* and $HAC1^{i}$*-HA* strains were obtained.

**Figure S3: Southern hybridization of *V. dahliae ODE1* deletion, and *ODE1-GFP* complementation strains.** *V. dahliae* wildtype (WT), *ODE1* deletion (Δ*ODE1* transformant number 12 and 16) and the complementation strain harbouring an *ODE1-GFP* construct under control of the native promoter and terminator at the endogenous locus (*ODE1-GFP* transformant number 1 and 2) were tested. Genomic DNA was cut with *Sca*I restriction enzyme and the 3´ flanking region of *ODE1* was used as a probe. $Hyg^{R}$: hygromycin resistance cassette under control of the *gpdA* promoter and a *trpC* terminator; $Nat^{R}$: nourseothricin resistance cassette under control of the *gpdA* promoter and a *trpC* terminator. **left:** Scheme of restriction sites (arrows) and expected fragment length. **right:** Predicted bands of 4.6 kb for wildtype, 3.6 kb for the deletion, and 2.9 kb for the endogenous complementation strains were observed.

**Figure S4: Phylogeny of Rok1-like phosphatases.** Relations of Rok1-like phosphatases are depicted in a phylogenetic tree with *Verticillium dahliae* (*VDAG_JR2_Chr7g08960a*), *Magnaporthe oryzae* (XP_003712767.1), *Neurospora crassa* (XP_962856.1), *Trichoderma reesei* (XP_006961240.1), *Aspergillus nidulans* (XP_662148.1), *Aspergillus fumigatus* (XP_749411.1), *Ustilago maydis* (UMAG_03701), *Saccharomyces cerevisiae* (NP_014345), *Homo sapiens* (NP_004408.1) sequences (ClustalW algorithm, scale bar = average number of amino acid substitutions per site). Rok1-like proteins with dual specificity phosphatase domains (IPR020422, yellow) are shown.

**Table S1: Verticillium strains constructed and used in this study.**

| Strain | Description | Reference |
|---|---|---|
| JR2/ WT | *Solanum lycopersicum* isolate | (Fradin *et al.*, 2009) |
| VGB45 | $^{p}gpdA:GFP:trpC^{t}:{}^{p}gpdA:HYG^{R}:trpC^{t}$ | (Bui *et al.*, 2019) |
| VGB279/ VGB280 | $\Delta HAM5::{}^{p}gpdA:NAT^{R}:trpC^{t}$ | This study |
| VGB331/ VGB332 | $\Delta ODE::{}^{p}gpdA:NAT^{R}:trpC^{t}$ | This study |
| VGB335/ VGB336 | $\Delta VMK1::{}^{p}gpdA:HYG^{R}:trpC^{t}$ | This study |
| VGB337/ VGB338 | $\Delta MEK2::{}^{p}gpdA:HYG^{R}:trpC^{t}$ | This study |
| VGB346/ VGB347 | $\Delta MEK2::{}^{p}gpdA:HYG^{R}:trpC^{t};$ $\Delta HAM5::{}^{p}gpdA:NAT^{R}:trpC^{t}$ | This study |
| VGB358/ VGB359 | $\Delta ODE1::{}^{p}gpdA:NAT^{R}:trpC^{t}::{}^{p}ODE1:ODE1:GFP:$ $^{p}gpdA:HYG^{R}:trpC^{t}:ODE1^{t}$ | This study |
| VGB371/ VGB372 | $\Delta HAC1::{}^{p}gpdA:HYG^{R}:trpC^{t}$ | This study |
| VGB380 | $\Delta HAC1::{}^{p}gpdA:HYG^{R}:trpC^{t};$ $^{p}gpdA:GFP:trpC^{t}:{}^{p}gpdA:NAT^{R}:trpC^{t}$ | This study |
| VGB382 | $\Delta HAC1::{}^{p}gpdA:HYG^{R}:trpC^{t};$ $^{p}HAC1:HAC1:HAC1^{t}:{}^{p}gpdA:NAT^{R}:trpC^{t}$ | This study |
| VGB388/ VGB389 | $\Delta MEK2::{}^{p}gpdA:HYG^{R}:trpC^{t};$ $^{p}MEK2:MEK2:MEK2^{t}:{}^{p}gpdA:NAT^{R}:trpC^{t}$ | This study |
| VGB392 | $^{p}gpdA:GFP:trpC^{t}:{}^{p}gpdA:NAT^{R}:trpC^{t}$ | This study |
| VGB413/ VGB414 | $\Delta VMK1::{}^{p}gpdA:HYG^{R}:trpC^{t};$ $^{p}VMK1:VMK1:VMK1^{t}:{}^{p}gpdA:NAT^{R}:trpC^{t}$ | This study |
| VGB415/ VGB416 | $\Delta HAM5::{}^{p}gpdA:NAT^{R}:trpC^{t};$ $^{p}HAM5:HAM5:HAM5^{t}:{}^{p}gpdA:HYG^{R}:trpC^{t}$ | This study |
| VGB417 | $\Delta MAK2::{}^{p}gpdA:HYG^{R}:trpC^{t};$ $\Delta HAM5::{}^{p}gpdA:NAT^{R}:trpC^{t}$ | This study |
| VGB439/ VGB440 | $\Delta HAC1::{}^{p}gpdA:HYG^{R}:trpC^{t};$ $^{p}HAC1: HAC1^{u}:HAC1^{t}:{}^{p}gpdA:NAT^{R}:trpC^{t}$ | This study |
| VGB437/ VGB438 | $\Delta HAC1::{}^{p}gpdA:HYG^{R}:trpC^{t};$ $^{p}HAC1: HAC1^{i}:HAC1^{t}:{}^{p}gpdA:NAT^{R}:trpC^{t}$ | This study |
| VGB477 | $^{p}gpdA:H2B:RFP:trpC^{t}:{}^{p}gpdA:GEN^{R}:trpC^{t}$ | This study |
| VGB493/ VGB494 | $\Delta ODE1::{}^{p}gpdA:NAT^{R}:trpC^{t}$ $^{p}gpdA:H2B:RFP:trpC^{t}:{}^{p}gpdA:GEN^{R}:trpC^{t}$ | This study |

$^{p}$: promoter, $^{t}$: terminator, $NAT^{R}$: nourseothricin resistance marker, $GEN^{R}$: geneticin resistance marker, $HYG^{R}$: hygromycin B resistance marker, two VGB numbers for one genotype indicate two independent transformants.

**Table S2: Primers used in this study.**

| Primer name | Primer sequence (5´→ 3´) | Length (bp) | Overhang to |
|---|---|---|---|
| JST76b | GTA TGT TGT GTG GAA AGC ACG GAG CAG AGA CCA | 33 | pPK2 |
| JST77a | GGT TCT GGT ACA CGA CGA GC | 20 | - |
| JST110 | GTA TGT TGT GTG GAA CCG CGA GGG TTG GAG AGG | 33 | pME4564 |
| JST111 | ACC GGT CAC TGT ACA GAC GGG CCT GAT ATT CTT TCG A | 37 | *^p^gpdA* |
| JST112 | AGG TAA TCC TTC TTT GTG GCC GTC TTT TCA CAG GC | 35 | *trpC^t^* |
| JST113 | CAC AGT ACA CGA GGA TCT CGT CCG GAC TGA TCC AA | 35 | pME4564 |
| JST127 | CGT ATG TTG TGTG GAA GGA TGG CCA ATG TGG ATT TGA T | 38 | pME4564 |
| JST128 | CAC CGG TCA CTG TAC AGG TAC TGG TGG CTC TTG GGA | 36 | *^p^gpdA* |
| JST129 | GAG GTA ATC CTT CTT TTC GGA TTG GAC AGT AGA CAA GTT TG | 41 | *^p^gpdA* |
| JST130 | CAC AGT ACA CGA GGA CGC GCA CAG TTA CAC TTC ATA CTC T | 40 | pME4564 |
| JST137 | TCC ACA TTG GCC ATC CTT CCA CAC AAC ATA CGA GCC G | 37 | *^p^ODE1* |
| JST138 | AGT GTA ACT GTG CGC GTC CTC GTG TAC TGT GTA AGC | 36 | *ODE1^t^* |
| JST171 | ATG GAG TCT TGG GAG CAC TC | 20 | - |
| JST172 | TCA GAC ACC AAC CGC AAT | 18 | - |
| JST174 | TCA GCG AAA GCG CAC TC | 17 | - |
| JST177 | AAA GAA GGA TTA CCT CTA AAC AAG TGT | 27 | - |
| JST178 | GGT ACC GAG CTC GAT TTA CTT GTA CAG CTC GTC CA | 35 | *^p^gpdA* |
| JST179 | ACC ACC GCT ACC ACC CTG CTC ATC CGT ACG GC | 32 | linker C-terminal *GFP* |
| JST180 | ATT CTT AAT TAA GAT GGA TGG CCA ATG TGG AT | 32 | pPK2 |
| JST184 | GTA TGT TGT GTG GAA CGA GTG GAG ATG TGG AGT | 33 | pME4815 |
| JST185 | ACC GGT CAC TGT ACA TGG CAT GCG GAG AGA C | 31 | pME4815 |
| JST186 | ATT CTT AAT TAA GAT GAC AAG AGT CAA GCC CAC | 33 | pME4564 |
| JST187 | AGA TCC CCG GGT ACC GAT GGA CGA AGC GAC TC | 32 | *^p^gpdA* |
| JST189 | AGG ACT TCT AGA AGG TCC AGC TCC AAA TCA ATT AAC C | 37 | pME4564 |

| Primer name | Primer sequence (5´→ 3´) | Length (bp) | Overhang to |
|---|---|---|---|
| JST211 | GGT CAC TGT ACA GAT GGG ACT CGT ACC ATG TTT C | 34 | *trpC*t |
| JST212 | TGT TGT GTG GAA GAT ACT AAG TAC TGG TTG TGG CTG AC | 38 | pME4815 |
| JST213 | GGT CAC TGT ACA GAT AGG CTT GGA GAT GAC GAG | 33 | p*gpdA* |
| JST216 | TGT TGT GTG GAA GAT GAC AAG AGT CAA GCC CAC | 33 | pME4564 |
| JST243 | TGT TGT GTG GAA GAT AGC ACG GAG CAG AGA CCA | 33 | pME4815 |
| JST244 | GGT CAC TGT ACA GAT CGA ACC GGT GAT GGA TAC G | 34 | p*gpdA* |
| JST245 | ATT CTT AAT TAA GAT CTG CTC CTA TTC GGC TCC | 33 | pPK2 |
| JST246 | GGT ACC GAG CTC GAT TCT CGT CCG GAC TGA TCC | 33 | pPK2 |
| JST266 | CTA TCC GCC GCT AGC GTA ATC GGG CAC ATC GTA TGG GTA GCC GCC GCT GAC ACC AAC CGC AAT GC | 65 | *HA* tag |
| JST267 | CTA TCC GCC GCT AGC GTA ATC GGG CAC ATC GTA TGG GTA GCC GCC GCT GCG AAA GCG CAC TCG T | 64 | *HA* tag |
| JST268 | CTA TCC GCC GCT AGC GTA | 18 | - |
| JST269 | TGT TGT GTG GAA GAT GCT GAG GTC ATG GCT GAC | 33 | pME4564 |
| JST270 | CTC CCA AGA CTC CAT TTT GGA CGG CTT TGT GTG | 33 | *HAC1* |
| JST271 | GCT AGC GGC GGA TAG GGG CTG TGA GAA TCG GGT | 33 | *HA* |
| JST272 | GGT CAC TGT ACA GAT GGG ACT CGT ACC ATG TTT CA | 35 | *trpC*t |
| JST273 | GCT AGC GGC GGA TAG TTT GAT TTT TAT CAT GAT GAC GG | 38 | *HA* |
| JST290 | TTC AGA AAA TCC TAA CCT GCA G | 22 | - |
| JST291 | ACA CCA ACC GCA ATG CCT | 18 | - |
| JS-V5 | CGT ATG TTG TGT GGA AAC TAA GTA CTG GTT GTG GCT GAC | 39 | pPK2 |
| JS-V6 | AAG ATC CCC GGG TAC CTT TGG GTG ATG TGC GTG G | 34 | p*gpdA* |
| JS-V7 | ACA ACC AGT ACT TAG TTT CCA CAC AAC ATA CGA G | 34 | p*MEK2* |
| JS-V8 | ACG CAC ATC ACC CAA AGG TAC CCG GGG ATC TTT C | 34 | p*MEK2* |
| JS-V9 | TCC TTC TTT CTA GAA GTT GAA CAG GCC TGT CTG G | 34 | pME4821 |

| Primer name | Primer sequence (5´→ 3´) | Length (bp) | Overhang to |
|---|---|---|---|
| JS-V10 | ACA CAG TAC ACG AGG AAG GCT TGG AGA TGA CGA G | 34 | pME482 |
| JS-V11 | CGT CAT CTC CAA GCC TTC CTC GTG TAC TGT GTA AG | 35 | *MEK2*[t] |
| JS-V12 | AGA CAG GCC TGT TCA ACT TCT AGA AAG AAG GAT TAC CTC | 39 | *MEK2*[t] |
| JS-V21 | GAG GTA ATC CTT CTT TGG TGG CAG TGG CAG TGG | 33 | pPK2 |
| JS-V22 | ACA CGA GGA CTT CTA GCG AAC CGG TGA TGG ATA CGT T | 37 | pPK2 |
| JS-V23 | ATC CAT CAC CGG TTC GCT AGA AGT CCT CGT GTA CTG T | *37* | *VMK1*[t] |
| JS-V24 | CAC TGC CAC TGC CAC CAA AGA AGG ATT ACC TCT AAA CAA GT | 41 | *VMK1*[t] |
| JT1 | CTC TAG AGG ATC CCC TGT ACA GTG ACC GGT GAC TC | 35 | pCOM |
| JT2 | TCG AGC TCG GTA CCC ACC TCT AAA CAA GTG TAC CTG T | 37 | pCOM |
| ML1 | TTC CAC ACA ACA TAC GAG CC | 20 | - |
| ML2 | TCC TCG TGT ACT GTG TAA GC | 20 | - |
| ML5 | TGT ACA GTG ACC GGT GAC TCT T | 22 | - |
| ML6 | TCC CGC GGT CGG CAT CTA CTT CAG GGG CAG GGC ATG CT | 38 | *trpC* terminator |
| ML7 | TGA GCA TGC CCT GCC CCT GAA GTA GAT GCC GAC CGC G | 37 | *NAT*[R] |
| ML8 | AAA GAA GGA TTA CCT CTA AAC AA | 23 | - |
| ML9 | TGT ACA GTG ACC GGT GAC | 18 | - |
| PC4 | TGT ACA GTG ACC GGT GAC TC | 20 | - |
| pJet1.2 reverse | AAG AAC ATC GAT TTT CCA TGG CAG | 24 | - |
| RH523 | ACG TCC TCG GAG GAG GCC ATG GTG ATG TCT GCT CAA GCG | 39 | *RFP* |
| RH524 | CCG CTT GAG CAG ACA TCA CCA TGG CCT CCT CCG AGG AC | 38 | [p]*gpdA* |
| RH525 | GGC ATA CCA CCG CTA CCA CCG GCG CCG GTG GAG TGG C | 37 | Linker, *H2B* |
| RH526 | GCG CCG GTG GTA GCG GTG GTA TGC CCC CCA AGG CCG C | 37 | Linker, *RFP* |
| RH527 | TCC CGC GGT CGG CAT CTA CTT TAT TTC GTG GAC GAG GAA TAC | 42 | *trpC*[t] |
| RH528 | ATT CCT CGT CCA CGA AAT AAA GTA GAT GCC GAC CGC GG | 38 | *H2B* |
| RH529 | TCT AGA AAG AAG GAT TAC CTC T | 22 | - |
| RH530 | AGT TCT AGA TGT ACA GTG ACC GGT GAC TC | 29 | Restriction site |

| Primer name | Primer sequence (5´→ 3´) | Length (bp) | Overhang to |
|---|---|---|---|
| RO3 | GGT ACC CGG GGA TCT TTC G | 19 | - |
| SAB16 | GGT GGT AGC GGT GGT ATG | 18 | - |
| SAB52 | GTA TGT TGT GTG GAA GAT ATC TGT ACA GTG ACC GGT GAC | 39 | pME4564 |
| SAB53 | CAC AGT ACA CGA GGA AAA GAA GGA TTA CCT CTA AAC AA | | pME4564 |
| SZ9 | AAC ACC CAG AAC AAG ATG CGC | 21 | - |
| SZ10 | GCT TGA CCT TGA GAT CCT TG | 20 | - |
| SZ11 | TGC ATT CTT GGC AAG AGA TGT GTG | 24 | - |
| SZ12 | AGC TTG TTA TCC TTG TCC TCG GT | 23 | - |

Blue: overhangs for fusion PCR, Seamless and FastCloning, or restriction sites, purple: overhang for fusion of linker or tag, $^{p}$: promoter, $^{t}$: terminator.

**Table S3: Plasmids constructed and used in this study.**

| Plasmid | Description | Reference |
|---|---|---|
| pCOM | $^{p}$*gpdA:GEN*$^{R}$*:trpC*$^{t}$*, KAN*$^{R}$*,* left and right border for ATMT | (Zhou *et al.*, 2013) |
| pGreen2 | $^{p}$*gpdA:HYG*$^{R}$*:trpC*$^{t}$; $^{p}$*gpdA:GFP:trpC*$^{t}$; *KAN*$^{R}$; left and right border for ATMT | (Tran *et al.*, 2014) |
| pJet1.2 | Cloning vector with *AMP*$^{R}$ | Thermo Fisher Scientific |
| pPK2 | Cloning vector with *KAN*$^{R}$ and *HYG*$^{R}$; left and right border for ATMT | (Covert *et al.*, 2001) |
| pKO2.0 | Cloning vector with *KAN*$^{R}$ and *NAT*$^{R}$: left and right border for ATMT | (Timpner *et al.*, 2013) |
| pME3857 | $^{p}$*gpdA:mRFP:H2A* | (Bayram *et al.*, 2012) |
| pME4564 | Cloning vector with *KAN*$^{R}$ and *HYG*$^{R}$; left and right border for ATMT | (Bui *et al.*, 2019) |
| pME4815 | $^{p}$*gpdA:NAT*$^{R}$*:trpC*$^{t}$ in pME4564, *KAN*$^{R}$; left and right border for ATMT | This study |
| pME4819 | $^{p}$*gpdA:GFP:trpC*$^{t}$ in pME4815; left and right border for ATMT | This study |
| pME4820 | Δ*HAM5::*$^{p}$*gpdA:NAT*$^{R}$*:trpC*$^{t}$ in pME4564; left and right border for ATMT | This study |
| pME4821 | $^{p}$*MEK2* in pPK2; left and right border for ATMT | This study |
| pME4822 | Δ*MEK2::*$^{p}$*gpdA:HYG*$^{R}$*:trpC*$^{t}$ in pPK2; left and right border for ATMT | This study |
| pME4823 | *HAM5*$^{t}$ in pPK2; left and right border for ATMT | This study |
| pME4824 | *VMK1*$^{t}$ in pPK2; left and right border for ATMT | This study |
| pME4825 | Δ*VMK1::*$^{p}$*gpdA:HYG*$^{R}$*:trpC*$^{t}$ in pPK2; left and right border for ATMT | This study |
| pME4826 | $^{p}$*MEK2:MEK2:MEK2*$^{t}$ in pME4815; left and right border for ATMT | This study |
| pME4827 | $^{p}$*VMK1:VMK1:VMK1*$^{t}$ in pME4815; left and right border for ATMT | This study |
| pME4828 | $^{p}$*HAM5:HAM5:HAM5*$^{t}$*:*$^{p}$*gpdA:HYG*$^{R}$*:trpC*$^{t}$ in pPK2; left and right border for ATMT | This study |
| pME4830 | Δ*HAC1::*$^{P}$*gpdA:HYG*$^{R}$*:trpC*$^{t}$ in pPK2, left and right border for ATMT | This study |
| pME4831 | $^{p}$*HAC1:HAC1:HAC1*$^{t}$ in pME4815, left and right border for ATMT | This study |
| pME4832 | *HAC1*$^{u}$*:HA* in pJet1.2 | This study |
| pME4833 | *HAC1*$^{i}$*:HA* in pJet1.2 | This study |
| pME4834 | $^{p}$*HAC1:HAC1*$^{i}$*:HAC1*$^{t}$ in pME4564, left and right border for ATMT | This study |

| Plasmid | Description | Reference |
|---|---|---|
| pME4835 | $^{p}$*HAC1:HAC1*$^{u}$*:HAC1*$^{t}$ in pME4564, left and right border for ATMT | This study |
| pME4836 | Δ*ODE1::*$^{p}$*gpdA:NAT*$^{R}$*:trpC*$^{t}$ in pME4564, left and right border for ATMT | This study |
| pME4837 | *ODE1*$^{t}$ in pPK2, left and right border for ATMT | This study |
| pME4838 | Δ*ODE1::*$^{p}$*gpdA:NAT*$^{R}$*:trpC*$^{t}$*::*$^{p}$*ODE1:ODE1:GFP:*$^{p}$*gpdA:HYG*$^{R}$*:trpC*$^{t}$*:ODE1*$^{t}$ in pME4837, left and right border for ATMT | This study |
| pME4973 | $^{p}$*gpdA:RFP:H2B:trpC*$^{t}$ in pJet1.2 | This study |
| pME4975 | $^{p}$*gpdA:RFP:H2B:trpC*$^{t}$ in pPK2, left and right border for ATMT | This study |
| pME4976 | $^{p}$*gpdA:RFP:H2B:trpC*$^{t}$ in pCOM, left and right border for ATMT | This study |
| pME4978 | $^{p}$*gpdA:NAT*$^{R}$*:trpC*$^{t}$ in pJet1.2, *KAN*$^{R}$ | This study |

*AMP*$^{R}$; ampicillin resistance marker, ATMT: *Agrobacterium tumefaciens* mediated transformation, *GEN*$^{R}$: geneticin resistance marker, *HYG*$^{R}$: hygromycin B resistance marker, *KAN*$^{R}$: kanamycin resistance marker, *NAT*$^{R}$: nourseothricin resistance marker, $^{p}$: promoter, $^{t}$: terminator

**Table S4: The cDNA sequence of *V. dahliae* JR2 *HAC1*$^{i}$.** The *V. dahliae* JR2 *HAC1*$^{i}$ sequence with 1254 nucleotides was obtained by RNA extraction and cDNA synthesis from mycelium of wildtype cultures grown in 50 ml SXM (1x10$^{7}$ spores) for 4 d under constant agitation at 25 °C and subsequent supplementation with 3 mM DTT for three hours. *HAC1*$^{i}$ was amplified using primers JST171 and JST174 and fully sequenced.

| | | |
|---|---|---|
| 1 | ATG GAG TCT TGG GAG CAC TCC ACC ACA CCA | 30 |
| 31 | ATG ATC AAG TTC GAG GAC TCG CCA GCC GAG | 60 |
| 61 | TCT TTC GTC TCG ACA CCA GGC GAC ATG TAC | 90 |
| 91 | CCG TCA CTC TTC CCA GAG TCC GCC TCC CCC | 120 |
| 121 | AAC ACC CTC GAT CCT TCC AAC ATG ATG AGC | 150 |
| 151 | CCT TCC TCA CCC CAA GAC CTC ACC ATT GCC | 180 |
| 181 | GAC ACG GAT ATG CCT CTC TCC GAG GCT TCC | 210 |
| 211 | GCC GGC GAC AAG AAG GGG TCC AAG AAG CGC | 240 |
| 241 | AAG TCC TGG GGT CAG GTC CTT CCC GAG CCC | 270 |
| 271 | AAG ACC AAC TTG CCG CCC AGG AAA CGA GCC | 300 |
| 301 | AAG ACT GAG GAT GAG AAG GAG CAG CGT CGT | 330 |
| 331 | GTG GAA CGC GTT CTG CGC AAC CGC CGT GCT | 360 |
| 361 | GCC CAG TCT TCG AGG GAG CGC AAG AGG CTC | 390 |
| 391 | GAG GTT GAG GCC CTC GAG ATG AAG AAC AAG | 420 |
| 421 | GAG CTC GAG ACT GCC CTG AAC CAC GCA CAA | 450 |
| 451 | CAG GCG AAC GCT AGG TTG ATG GAG GAG CTT | 480 |
| 481 | ACC AAG TTC CGC CGT GGT TCC GGT GCC GTC | 510 |
| 511 | GCC CGT TCT TCT TCC CCC TTT GAC TCC TTC | 540 |
| 541 | CAC AAC AGC AAC TCG GTC ACC CTC TCC CCC | 570 |
| 571 | GAG CTG TTC GGC TCT CAA GAC GGC CGC CGG | 600 |
| 601 | CCA TCA GTG GCC GAC TCC GAG TCG ACA CTC | 630 |
| 631 | GTC GAC GGT TTG ATG GCG GCC TCC AAG TCC | 660 |
| 661 | GCC GCG ACC GTC AAC CCC GCC TCC CTC TCG | 690 |
| 691 | CCC GCC CTC ACC CCC GTC CCC GAG ACG GAT | 720 |
| 721 | GAG ACC AGC GCC CAA CAA GAA GCT GCC GTG | 750 |
| 751 | GCC GCC CCT TCC CCT GTC GCC CTT TCC TCC | 780 |
| 781 | GAC GTG ACA CAA CGT CCT GCC GTG TCG GTC | 810 |
| 811 | GGA GGA AAT GCC TCA GTC GTG GGT GGC CTC | 840 |
| 841 | GCA GAC TTC CCT GCA CCC AAC ATG GAC TTT | 870 |
| 871 | GTA CCT TCA GCT TCA GAT GCT CAT GAT CAC | 900 |
| 901 | TTC CTC GGC GGT CAT TTC AGC GTG TCA GAG | 930 |
| 931 | GCC TTT GAT GCA GAT CGC TAT GTC CTT GAG | 960 |
| 961 | AGC GGG CTT CTC TCT TCC CCC AAC TCA GTC | 990 |
| 991 | GAT TAT GAC AAC GAT ATT ATG GCT GGT GAC | 1020 |
| 1021 | TCG TCC GCG TTC GCA TCC GCG TTC AAC TTC | 1050 |
| 1051 | GAC ATG GAC GAG TTC CTC AAC GAT GAG GCC | 1080 |
| 1081 | AGC GCA GCC GCC ACT GAC GCG TCA GCA GCG | 1110 |
| 1111 | GAG AAC AGC GCA GCG GAC CCG GAC TAC GGC | 1140 |
| 1141 | CGC CGT GCC CTT AAC CCT GAG ACT CAA GTC | 1170 |
| 1171 | TCT TCA GAA AAT CCT AAC CTG CAG CCC CAA | 1200 |
| 1201 | TCT GGC GCG TCC ACT TAT GGA TGC GAC GAT | 1230 |
| 1231 | GGA GGC ATT GCG GTT GGT GTC TGA | 1254 |

**Table S5: The amino acid sequence of *V. dahliae* JR2 Hac1.** The deduced protein sequence from *HAC1*$^{i}$ is 417 aa in length. Red: NLS predicted by cNLS Mapper (94-105 aa); Blue: N-terminal basic-leucine zipper domain (bZIP, PS50217; 107-164 aa).

| | | | | | |
|---|---|---|---|---|---|
| 1 | MESWEHSTTP | MIKFEDSPAE | SFVSTPGDMY | PSLFPESASP | 40 |
| 41 | NTLDPSNMMS | PSSPQDLTIA | DTDMPLSEAS | AGDKKGSKKR | 80 |
| 81 | KSWGQVLPEP | KTNLPPRKRA | KTEDEKEQRR | VERVLRNRRA | 120 |
| 121 | AQSSRERKRL | EVEALEMKNK | ELETALNHAQ | QANARLMEEL | 160 |
| 161 | TKFRRGSGAV | ARSSSPFDSF | HNSNSVTLSP | ELFGSQDGRR | 200 |
| 201 | PSVADSESTL | VDGLMAASKS | AATVNPASLS | PALTPVPETD | 240 |
| 241 | ETSAQQEAAV | AAPSPVALSS | DVTQRPAVSV | GGNASVVGGL | 280 |
| 281 | ADFPAPNMDF | VPSASDAHDH | FLGGHFSVSE | AFDADRYVLE | 320 |
| 321 | SGLLSSPNSV | DYDNDIMAGD | SSAFASAFNF | DMDEFLNDEA | 360 |
| 361 | SAAATDASAA | ENSAADPDYG | RRALNPETQV | SSENPNLQPQ | 400 |
| 401 | SGASTYGCDD | GGIAVGV | | | |

### Methods S1

#### Bacterial and fungal strains and cultivation conditions

*Escherichia coli DH5α* cells (Invitrogen) and *Agrobacterium tumefaciens* AGL-1 strain (Lazo *et al.*, 1991) were used for plasmid preparation and Verticillium transformation, respectively. *E. coli* was cultivated at 37 °C in lysogeny broth (LB) (Bertani, 1951) or on LB plates supplemented with kanamycin (100 µg/ml, AppliChem) or ampicillin (100 mg/ml, Carl Roth). *A. tumefaciens* was cultivated in LB medium supplemented with kanamycin (100 µg/ml) at 25 °C or 28 °C, respectively.

*V. dahliae* conidia were used for inoculation in liquid simulated xylem medium (SXM) (modified from Neumann & Dobinson, 2003 as described in (Hollensteiner *et al.*, 2017)) for spore production and potato dextrose bouillon (Carl Roth) for mycelium production and incubated at 25 °C shaking at 120 rpm. Conidia were harvested from SXM cultures after 5 to 7 d of incubation with sterile Miracloth (Calbiochem, Merck) and resuspended in sterile water. Conidiospore concentrations were determined using the Coulter Z2 Particle Count and Size Analyzer (Beckman Coulter) and the appropriate Coulter Isoton II Diluent. Mycelium was harvested after 4 to 6 d through Miracloth, rinsed with 0.96 % NaCl solution, dried with tissues and frozen in liquid nitrogen prior to isolation of genomic DNA, RNA or proteins.

Verticillium transformants were selected on potato dextrose medium (PDM, "Potato dextrose agar" (Carl Roth) with 0.5% additional agar) supplemented with nourseothricin (72 µg/ml; clonNAT, Werner BioAgents) or hygromycin B (50 µg/ml, InvivoGen), and cefotaxime (300 µg/ml; Wako chemicals) to eliminate *A. tumefaciens* cells.

PDM, SXM, modified Czapek Dox medium (CDM, 3% (w/v) sucrose, 2% (v/v) 50x AspA (3.5 mM $NaNO_3$, 350 mM KCl, 550 mM $KH_2PO_4$, pH 5.5), 2 mM $MgSO_4$, 1 ml of 1% (w/v) $FeSO_4$ solution and 2% (w/v) agar) (Smith, 1949), CDM with 3% (w/v) cellulose, 3% (w/v) galactose, or 3% (w/v) glucose as alternative carbon sources, CDM supplemented with 0.125% linoleic acid (LA), and CDM supplemented with different stress inducing agents (0.00075% $H_2O_2$, 0.004% SDS, 0.5 M NaCl, 0.8 M Sorbitol, 1 µg/ml tunicamycin) were used for analysis of fungal growth and *ex planta* phenotypes. Plates were incubated at 25 °C and growth was observed at indicated time points.

**DNA manipulation and strain construction**

The gene predictions for *V. dahliae* JR2 *HAM5* (*VDAG_JR2_Chr4g07170a*), *MEK2* (*VDAG_JR2_Chr1g13070a*), *VMK1* (*VDAG_JR2_Chr2g01260a*), and *ODE1* (*VDAG_JR2_Chr1g29610a*) were used from Ensemble Fungi (https://fungi.ensembl.org) and were confirmed on cDNA level. The transcript variant given for *HAC1* (*VDAG_JR2_Chr2g09780a*) was confirmed as the uninduced variant *HAC1ᵘ*. A second splice variant of the *HAC1* mRNA named *HAC1ⁱ* was identified (sequence given in Supporting Information Table S4). The cDNA used for amplification of *HAC1* was generated from RNA isolated from wildtype cultures grown in 50 ml SXM ($1x10^7$ spores) and incubated at 25°C under constant agitation for 4 d for the uninduced *HAC1* mRNA variant, and with subsequent supplementation with 3 mM DTT for 3 h for the induced *HAC1* mRNA variant. The reverse transcribed 1581 bp uninduced *HAC1* mRNA sequence was amplified with primers JST171 and JST172. The 1254 bp induced splice variant of *HAC1* was amplified using primers JST171 and JST174. Transcripts were fully sequenced.

All fragments were PCR amplified using the Q5 Hot Start polymerase (NewEnglandBiolabs) or Phusion polymerase (Thermo Scientific). Fungal transformants were verified by Southern hybridization (Bui *et al.*, 2019).

For construction of deletion and complementation cassettes the GeneArt Seamless Cloning and Assembly Kit (Thermo Scientific) and the FastCloning protocol (Li *et al.*, 2011) was used. Therefore 1-2 kb up- and downstream flanking sequences were amplified from *V. dahliae* JR2 genomic DNA with 15-16 bp homologous overhangs to the desired neighbouring fragments. The PCR products were purified with NucleoSpin Gel and PCR Clean-up Kit (Macherey-Nagel) and used for transformation of *E. coli* DH5α. All plasmids were verified by Sanger sequencing performed by the Microsynth Seqlab in Göttingen.

The nourseothricin resistance marker cassette used for the generation of knock-out strains in this study was constructed by fusion PCR and cloned into the pJet1.2 Cloning vector, resulting in pME4978. The *gpdA* promoter and the nourseothricin resistance marker gene were amplified from vector pKO2.0 using the primers ML5 and ML6. The *trpC* terminator was amplified from pPK2 using primers ML7 and ML8.

The pME4564 vector was amplified using the primers ML1 and ML2. The nourseothricin marker cassette, consisting of the *gpdA* promoter fused to the *NAT*$^R$ marker fused to a *trpC* terminator was amplified from pME4978 with the primers

SAB52 and SAB53 and fused to the pME4564 backbone via Seamless cloning, resulting in pME4815.

Plasmid and strain construction of the *HAM5* single and double deletion with *VMK1*

For construction of the *HAM5* (*VDAG_JR2_Chr4g07170a*) deletion cassette the 1500 bp flanking region 333 bp upstream of the ORF was amplified with primers JST110 and JST111, and the 1000 bp downstream flanking region was amplified with primers JST112 and JST113 from wildtype fungal DNA. The 2194 bp $NAT^R$ marker cassette was amplified with ML8 and ML9 from pME4815. The fragments were ligated into the 6728 bp backbone pME4564 amplified with primers ML1 and ML2. The resulting plasmid pME4820 was used for wildtype and Δ*VMK1* (VGB335) transformation. 5´ and 3´ flanking regions of *HAM5* were amplified with primers mentioned before and labelled as probes for Southern hybridization. Restriction enzymes *Xho*I (5´ flanking region) and *Sac*I (3´ flanking region) were used to cut genomic DNA. The resulting *HAM5* single deletion transformants were conserved as VGB279 and VGB280, the *HAM5* and *VMK1* double deletion transformant was conserved as VGB417.

Plasmid and strain construction of the ectopic *HAM5* complementation

For construction of the ectopic *HAM5* complementation cassette a 7265 bp sequence including 1359 bp 5´ flanking region, 4906 bp *HAM5* ORF, and 1000 bp 3´ flanking region was amplified using JST245 and JST246 from fungal genomic DNA. The PCR product was ligated into pPK2 harbouring the $HYG^R$ marker cassette cut with the restriction enzyme *Eco*RV. The resulting plasmid was named pME4828 and used for Δ*HAM5* transformation. Flanking regions of *HAM5* were amplified using primers mentioned before and labelled as probes for Southern hybridization. Restriction enzymes *Pvu*II (5´ flanking region) and *Sac*I (3´ flanking region) were used to cut genomic DNA. The resulting *HAM5-C* complementation transformants were conserved as VGB415 and VGB416.

Plasmid and strain construction of the *VMK1* deletion

For construction of the *VMK1* (*VDAG_JR2_Chr2g01260a*) deletion cassette the 898 bp 3´ flanking region was amplified with primers JS-V21 and JS-V22 from fungal wildtype DNA and ligated to the 10739 bp pPK2 backbone with $HYG^R$ marker cassette amplified with JS-V23 and JS-V24, resulting in pME4824. The 1472 bp 5´ flanking region was amplified with primers JST77a and JST76b from fungal wildtype DNA and cloned into pME4824, amplified with primers ML1 and JS-V23, resulting in pME4825 used for wildtype transformation. Flanking regions of *VMK1* were amplified using primers JST77a and JST76b (5´ flanking region) and JS-V21 and JS-V22 (3´ flanking region) and labelled as probes for Southern hybridization. Restriction enzymes *Xho*I (5´) or *Bgl*I (3´) were used to cut genomic DNA. The resulting *VMK1* deletion transformants were conserved as VGB335 and VGB336.

Plasmid and strain construction of the ectopic *VMK1* complementation

For construction of the ectopic *VMK1* complementation cassette a 3661 bp sequence, including 1473 bp 5´ flanking region, 1260 bp *VMK1* ORF, and 928 bp 3´ flanking region, was amplified from fungal wildtype DNA using JST243 and JST244. The PCR product was ligated to pME4815 harbouring a $NAT^R$ marker cassette cut with the restriction enzyme *Eco*RV, resulting in the plasmid pME4827 used for Δ*VMK1* transformation. Flanking regions of *VMK1* were amplified using the same primers as for Southern hybridization of the *VMK1* deletion strain and labelled as probes for Southern hybridization. The restriction enzymes *Vsp*I (5´ flanking region) or *Bgl*I (3´ flanking region) were used to cut genomic DNA. The resulting *VMK1-C* complementation transformants were conserved as VGB413 and VGB414.

Plasmid and strain construction of the *MEK2* single and double deletion with *HAM5*

For construction of the *MEK2* (*VDAG_JR2_Chr1g13070a*) deletion cassette the 1500 bp 5´ flanking region was amplified with primers JS-V5 and JS-V6 from fungal wildtype DNA and ligated to the 10676 bp pPK2 backbone with $Hyg^R$ marker cassette amplified with JS-V7 and JS-V8, resulting in pME4822.

The 1500 bp 3´ flanking region 385 bp downstream of the ORF was amplified with primers JS-V9 and JS-V10 from fungal wildtype DNA and ligated to pME4822 amplified with primers JS-V11 and JS-V12, resulting in pME4822 used for wildtype and Δ*HAM5* (VGB279) transformation. Flanking regions of *MEK2* were amplified using the primers mentioned before and labelled as probes for Southern hybridization. The restriction enzymes *Hind*III (5´ flanking region) or *Nru*I (3´ flanking region) were used to cut genomic DNA. The resulting *MEK2* single deletion transformants were conserved as VGB337 and VGB338, the *MEK2* and *HAM5* double deletion transformants were conserved as VGB346 and VGB347.

Plasmid and strain construction of the ectopic *MEK2* complementation

For construction of the ectopic *MEK2* complementation cassette a 5176 bp sequence including 1500 bp 5´ flanking region, 1756 bp *MEK2* ORF, and 1885 bp 3´ flanking region was amplified from fungal wildtype DNA using JST212 and JST213. The PCR product was ligated to pME4815 harbouring a *NAT*$^R$ marker cassette cut with the restriction enzyme *EcoR*V, resulting in the plasmid pME4826 used for Δ*MEK2* transformation. Flanking regions of *MEK2* were amplified using the primers mentioned before and labelled as probes for Southern hybridization. The restriction enzymes *Hind*III (5´ flanking region) or *Nru*I (3´ flanking region) were used to cut genomic DNA. The resulting *MEK2-C* complementation transformants were conserved as VGB388 and VGB389.

<u>Plasmid and strain construction of the *HAC1* deletion</u>

For construction of the *HAC1* (*VDAG_JR2_Chr2g09780a*) deletion cassette a 1500 bp flanking region 218 bp upstream from the ORF was amplified using JST186 and JST187, and a 1445 bp 3´ flanking region was amplified using JST188 and JST189 from fungal wildtype DNA. The 3942 bp *Hyg*$^R$ marker cassette was amplified from pPK2 with primers ML8 and RO3. The fragments were ligated to pME4564 cut with restriction enzymes *EcoR*V and *Stu*I, resulting in pME4830 used for wildtype transformation. 1500 bp up- and 300 bp downstream flanking regions of *HAC1* were amplified using primers JST186 and JST187 (5´ flanking region) and JST272 and JST273 (3´ flanking region) and labelled as probes for Southern hybridization. Restriction enzymes *Sal*I (5´ flanking region) and *Sal*I (3´ flanking region) were used to cut genomic DNA. The resulting *HAC1* deletion transformants were conserved as VGB371 and VGB372.

<u>Plasmid and strain construction of the ectopic *HAC1* complementation</u>

For construction of the ectopic *HAC1* complementation cassette a 3636 bp insert containing 1702 bp 5´ flanking, 1634 bp *HAC1* gene, and 300 bp 3´ flanking region was amplified with primers JST216 and JST211 and cloned into pME4815 cut with the restriction enzyme *EcoR*V, resulting in plasmid pME4831 used for transformation of the Δ*HAC1* strain.

For Southern hybridization the same probes and restriction enzymes as for the *HAC1* deletion strain were used. The resulting *HAC1-C* complementation transformant was conserved as VGB382.

<u>Plasmid and strain construction of the ectopic *HAC1*$^u$*-HA* complementation</u>

For construction of a *HAC1*$^u$*-HA* complementation construct the uninduced mRNA splice variant of *HAC1* without stop codon was amplified with primers JST171 and JST172 from cDNA isolated from wildtype cultures incubated in SXM for 4 d. The C-terminal *HA*-tag was fused to *HAC1*$^u$ via amplification with a 48 bp *HA* sequence as overhang using JST171 and JST267. *HAC1^u-HA* was ligated to the cloning vector pJet1.2, resulting in pME4832.

The 1400 bp 5´ flanking region was amplified with primers JST269 and JST270, and a 300 bp 3´ flanking region was amplified with primers JST272 and JST273 from fungal wildtype DNA. The 1626 bp *HAC1^u-HA* sequence was amplified from pME4832 with JST171 and JST268. The fragments were ligated to pME4815 cut with the restriction enzyme *Eco*RV, resulting in pME4834, used for transformation of the Δ*HAC1* strain. For Southern hybridization the same probes as for the *HAC1* deletion strain were used. The restriction enzymes *Pvu*II (5´ flanking region) and *Sal*I (3´ flanking region) were used to cut genomic DNA. The resulting *HAC1^u-HA* transformants were conserved as VGB439 and VGB440.

<u>Plasmid and strain construction of the ectopic *HAC1^i-HA* complementation</u>

For construction of a *HAC1^i-HA* complementation construct the induced splice variant of *HAC1* without stop codon was amplified from cDNA, isolated from wildtype cultures incubated in SXM for 4 d with subsequent supplementation with 3 mM dithiothreitol (DTT) for 3 h using primers JST171 and JST174. The C-terminal *HA*-tag was fused to *HAC1^i* via amplification with a 48 bp *HA* sequence as overhang using JST171 and JST266. *HAC1^i-HA* was ligated to the cloning vector pJet1.2, resulting in pME4833 used for transformation of the Δ*HAC1* strain. The 1400 bp 5´ flanking region was amplified with primers JST269 and JST270, and a 607 bp 3´ flanking region was amplified with primers JST271 and JST272 from fungal wildtype DNA. The 1299 bp *HAC1^i-HA* sequence was amplified from pME4833 with JST171 and JST268. The fragments were ligated to pME4815 cut with the restriction enzyme *Eco*RV, resulting in pME4835 used for transformation of the Δ*HAC1* strain.

For Southern hybridization the same probes and restriction enzymes as for the *HAC1-C* strain were used. The resulting *HAC1^i-HA* transformants were conserved as VGB437 and VGB438.

<u>Plasmid and strain construction of the ectopic *GFP* overexpression strains WT *OE-GFP^NAT* and *ΔHAC1 OE-GFP*</u>

For construction of the *GFP* overexpression vector with *NAT^R* resistance marker cassette a 2378 bp fragment containing *GFP* under control of the *gpdA* promoter and *trpC* terminator was amplified from pGreen2 with primers JST184 and JST185 and ligated to pME4815, amplified with primers ML1 and ML9, resulting in pME4819 used for wildtype and Δ*HAC1* strain transformation.

The resulting WT *OE-GFP*$^{NAT}$ transformant was confirmed by fluorescence microscopy and phenotypic comparison to wildtype and conserved as VGB392.

The resulting Δ*HAC1 OE-GFP* transformant was confirmed by fluorescence microscopy, phenotypic comparison to the Δ*HAC1* strain and Southern hybridization. The 300 bp downstream flanking region of *HAC1* was amplified using primers JST272 and JST273 and labelled as probe for Southern hybridization. The restriction enzyme *Sal*I (3´) was used to cut genomic DNA. The resulting Δ*HAC1 OE-GFP* transformant was conserved as VGB380.

<u>Plasmid and strain construction of the *ODE1* deletion</u>

For construction of the *ODE1* (*VDAG_JR2_Chr1g29610a*) deletion cassette the 1000 bp 5´ and 1522 bp 3´ flanking regions were amplified from fungal wildtype DNA with primers JST127 and JST128 (5´) or JST129 and JST130 (3´). The 2194 bp *NAT*$^{R}$ marker cassette was amplified with ML8 and ML9 from pME4815 and ligated to the 6728 bp pME4564 backbone amplified with JST137 and JST138, resulting in pME4836 used for wildtype transformation. Flanking regions of *ODE1* were amplified using the primers mentioned before and labelled as probes for Southern hybridization. The restriction enzymes *Bgl*I (5´) or *Sca*I (3´) were used to cut genomic DNA. The Δ*ODE1* strains were conserved as VGB331 and VGB332.

<u>Plasmid and strain construction of endogenous C-terminally *GFP*-tagged *ODE1* complementation</u>

For construction of the *ODE1* complementation cassettes with either N- or C-terminal *GFP*-tag the 1522 bp 3´ flanking region was amplified from fungal wildtype DNA with primers JST129 and JST130 and ligated to the 10744 bp pPK2 backbone amplified with JST138 and JST177, resulting in plasmid pME4837.

For construction of the endogenous *ODE1-GFP* complementation cassette a 2501 bp PCR product containing the 1000 bp 5´ flanking region and 1501 bp *ODE1* gene without stop codon was amplified from fungal wildtype DNA with JST179 and JST180. The sequence of 720 bp C-terminal *GFP* with a 15 bp linker was amplified with primers SAB16 and JST178 from pGreen2. Both inserts were ligated to pME4837 cut with the restriction enzyme *EcoR*V, resulting in plasmid pME4838 used for transformation of the Δ*ODE1* strain.

Flanking regions of *ODE1* were amplified using the primers mentioned before and labelled as probes for Southern hybridization. The restriction enzymes *Ava*II (5´ flanking region) or *Sca*I (3´ flanking region) were used to cut genomic DNA.

The expression of the fusion protein was confirmed by fluorescence microscopy and immunoblotting with a GFP antibody. The strains were conserved as VGB358 and VGB359 (*ODE1-GFP*), and VGB360 and VGB361 (*GFP-ODE1*).

Plasmid and strain construction of WT *Histone-RFP* and *ODE1-GFP Histone-RFP* strains

For visualization of nuclei, a plasmid with RFP fused to H2B (*VDAG_JR2_Chr2g01720a*) under the control of the *gdpA* promotor was generated. The *gpdA* promotor was amplified from pKO2.0 using the primers PC4 and RH523. The RFP fragment was generated from pME3857 with the primers RH524 and RH525. Genomic DNA from the wildtype JR2 served as template for *H2B* and the primers RH526 and RH527 were used. The *trpC* terminator was amplified from pPK2 using the primers RH528 and RH529. The single fragments were fused by fusion PCR using the primers PC4 and RH529. The fusion product was ligated into the pJet1.2 vector, resulting in pME4973. The *RFP-H2B* expression cassette was amplified from pME4973 using the primers pJet1.2 reverse and RH530. The fragment as well as the vector pPK2 was restricted with *Xba*I and ligated, resulting in pME4975. To generate a plasmid with genetecin marker, the *RFP-H2B* expression cassette was amplified from pME4975 using the primers JT1 and JT2. The plasmid pCOM was linearized with *Sma*I. The *RFP-H2B* expression cassette was inserted into the vector using the Seamless cloning and assembly kit, resulting in pME4976. pME4976 was used for transformation of the wildtype JR2 and VGB358, resulting in VGB477, and VGB493/VGB494.

**Growth test**

Growth tests of Verticillium strains were performed by spotting of $5x10^4$ conidiospores on indicated media. Colony diameters were determined by measurement of two perpendicular diameters per colony. Growth was quantified by measurement of two perpendicular diameters per colony for three plates per transformant and medium (≙n=1). Significances were calculated using the one-way Anova and Student´s t-test from *SISA* online tool. Colony cross sections were analysed with a binocular microscope SZX12-ILLB2-200 (Olympus Deutschland) and microsclerotia were observed with an Axiolab light microscope (ZEISS).

**Quantification of colony centre melanization**

For quantification of melanization $5x10^4$ freshly harvested spores were spot inoculated on three 30 ml CDM with cellulose medium plates per transformant. Pictures from colonies were taken 9 d after spot inoculation from the top view after removal of aerial mycelium and the melanized area was measured from eight bit greyscale pictures using ImageJ software (Rasband, 1997). The means of the brightness factor were determined, set relative to wildtype and inverted. The mean value of three colonies per transformant was considered as one biological replicate (≙n=1). Significances were calculated using the one-way Anova and Student´s t-test from *SISA* online tool.

**Conidiospore quantification**

For spore quantification freshly harvested conidia were inoculated in liquid SXM to a concentration of $4x10^3$ conidiospores per ml in triplicates and incubated for 5 d under constant agitation at 135 rpm in four independent experiments. Spores were harvested through Miracloth (Calbiochem Merck), diluted in equal volumes of sterile water and conidiospore numbers were determined and normalized to wildtype. The values determined for two independent *ΔHAC1* transformants were summed up in one bar. Error bars indicate the standard deviations. Significances were calculated using the one-way Anova and Student´s t-test from *SISA* online tool.

**RNA extraction and cDNA synthesis**

RNA was purified using the Direct-zol RNA MiniPrep Kit (Zymo Research) according to the manufacturer´s instructions. Approximately 1 ml of mycelium powder was mixed with 1 ml TRIzol (Ambion and life technologies), and frozen in liquid nitrogen. DNase I was used to cut remaining DNA on columns according to the manufacturer´s protocol. Purified RNA was eluted in prewarmed DNase/RNase free water. Reverse transcription of 0.8 µg RNA to cDNA was performed using the QuantiTect Reverse Transcription Kit (Qiagen) according to the manufacturer´s protocol.

**Fungal genomic DNA purification**

The extraction method was modified from Kolar *et al.*, 1988. Mycelium was rinsed with 0.96% NaCl solution, dried, frozen and ground to fine powder in liquid nitrogen. The powder was mixed with 800 µl of lysis buffer [50 mM Tris pH 7.5, 50 mM EDTA pH 8, 3% (w/v) SDS and 1% (v/v) ß-mercaptoethanol], incubated at 65°C for one hour, and mixed with 800 µl phenol. The mixture was centrifuged for 20 min at 13000 rpm and 4°C. The upper phase was transferred into a new tube, mixed with 500 µl chloroform and centrifuged for 10 min at 13000 rpm and 4°C. The upper phase was transferred into a new tube, mixed with 400 µl isopropanol and centrifuged for two minutes at 13000 rpm. The precipitated genomic DNA was washed with 300 µl 70% ethanol and centrifuged for one minute at 13000 rpm. Ethanol was removed and the sediment was dried at 65°C for 20 min. DNA was resuspended in deionized $H_2O$ and RNA was cut by RNase A (10 mg/ml) at 65°C for 30 min.

**Quantification of gene expressions**

Transcription was analysed by qRT PCR. Histone *H2A* (*VDAG_JR2_Chr4g01430a*) and *EIF2B (VDAG_JR2_Chr4g00410a*) served as reference genes. Transcription levels were analysed in triplicates using a CFX Connect Real Time System cycler (Biorad) with Mesa Green qPCR MasterMix Plus for SYBR Assay (Eurogentec) and cycling was 2:20 min 95°C followed by 40 cycles of: 95 °C for 20 s, 60 °C for 22 s, and 72 °C for 22 s. Specificity of PCR products was tested with the subsequent melting curve analysis from 65 °C to 95 °C with 5 s per 0.5 °C after 10 s at 95 °C. Primers JST290 and JST291 (*HAC1* variants), SZ9 and SZ10 (*H2A*), and SZ11 and SZ12 (*EIF2B*) were used. Expression levels were quantified relative to reference genes in ΔΔCT method (Livak & Schmittgen, 2001). Significances were calculated using the one-way Anova and Student´s t-test from *SISA* online tool.

**Protein extraction and western hybridization**

Protein extracts were obtained from mycelial powder mixed with B* buffer (300 mM NaCl, 100 mM Tris-HCl pH 7.5, 10% glycerol, 2 mM EDTA, 0.02% $NP_4O$), 2 mM DTT, and cOmplete Protease inhibitor cocktail mix (Roche). The mixture was centrifuged for 30 min at 13000 rpm and 4 °C. The concentration of proteins was determined by Bradford assay (Bradford, 1976) and absorbance of the protein sample was determined with Roti-Quant solution (Carl Roth) and the Infinite M200 microplate reader operated with Magellan software (Tecan Trading) with a bovine serum albumin (BSA; Carl Roth) dilution series as standard. 80 µg protein extracts were loaded on 12 % SDS gels and blotted onto nitrocellulose membranes (Amersham Protran 0.45 µm, GE Healtcare Life Sciences). Ponceau S (0.2% Ponceau S, 3% TCA) staining was used for visualization of transferred proteins as loading control.

Membranes were incubated in mouse monoclonal α-GFP (Santa Cruz Biotechnology) or α-HA (1:2000; Sigma-Aldrich Chemie). As secondary antibody α-mouse 115-035-003 (Jackson Immuno Research) was used. Detection of proteins was conducted via horseradish peroxidase (HRP) substrate luminol based chemiluminescence and signals were detected with the Fusion SL chemiluminescence detector (Peqlab Biotechnolog), operated with the corresponding software Fusion 15.18 (Vilber Lourmat), and Amersham Hyperfilm ECL film (GE Healtcare Life Sciences), which was developed by the use of the Optimax (Protec) film processor.

**_Arabidopsis thaliana_ root colonization assay**

A Verticillium root colonization assay was performed with *A. thaliana* (Col-0) plants. The root colonization assay was modified from (Tran *et al.*, 2014). The *A. thaliana* seeds were surface sterilized for five minutes in a solution containing 70% (v/v) EtOH and 0.05% (v/v) Tween80 and dried at 35°C for 20 min. The sterilized seeds were placed on Murashige and Skoog (Murashige & Skoog, 1962) plates [0.22% MS + vitamin (Duchefa), 0.05% MES monohydrate (Carl Roth), 1% sucrose and 1.5% plant agar (Duchefa), pH 5.7], incubated at 4 °C overnight and transferred to the climate chamber at long day conditions with 16 h of light (fluorescence: 60, GroLEDs: 100, illumination: 95 µmol) at 25°C and eight hours of darkness at 22 °C. After 21 d plants were transferred to 1% water agarose. One day later roots were inoculated in fresh conidial suspensions ($1x10^5$ spores/ml) for 35 min and transferred back to 1% water agarose. Two third of the plate were covered with aluminium foil to shade the roots. Afterwards plants were incubated in a plant chamber at long day conditions for the indicated time.

For microscopy roots were cut and incubated for five minutes in the dark in a staining solution [0.0025% (v/v) propidium iodide and 0.005% (v/v) silwet]. The roots were transferred to an object slide with 150 µl $H_2O$ and 50 µl staining solution. Fungal root colonization was examined by fluorescence microscopy. Overview pictures [20x objective (Zeiss)] as well as close up pictures [63x objective (Zeiss)] were taken with 300 ms exposure time for GFP signals and 800 ms exposure time for RFP signals. Per experiment, fungal strain and time point root colonization was analysed for two to three independent plants.

**Plant infection experiments**

Per experiment 15 ten-day-old seedlings of *Solanum lycopersicum* (moneymaker) (Bruno Nebelung Kiepenkerl- Pflanzenzüchtung) were uprooted, wounded by striking and infected with $1x10^7$ conidiospores/ml by root dipping into 50 ml conidiospore solution or deionized sterile water as control for 40 min on a shaker at ~30 rpm. The seedlings were transferred to pots containing a sand:soil (1:1) mixture. The soil was inoculated with $3x10^7$ conidiospores or sterile water for mock plants per pot. The plants were allowed to grow in a BrightBoy GroBank climate chamber under long day conditions. The disease symptoms were scored at 21 d post infection according to the following disease rating criteria: The fresh weight excluding the roots, the length of the longest leaf, and the height of the vegetation point per plant were determined and

transformed into a score ranging from 1 to 5 relative to the mean values determined for the uninfected Mock plants (height/ length of the longest leave/ weight scores: 100-70% (Mock) = 1; 69-60% (Mock) = 2; 59-40% (Mock) = 3; <40% (Mock) = 4; dead plant = 5). The disease score per plant was calculated by the mean of the scores for each parameter (height/ length of the longest leave/ weight) ranging from 1 to 5 (1 = healthy, 2 = weak symptoms, 3 = strong symptoms, 4 = very strong symptoms, 5 = dead plant). The amount of plants categorized in the disease scores 1 to 5 relative to the total amount of tested plants was visualized in a stack diagram. Plants inoculated with spores of two independent transformants with the same genotype were added up in one bar. Student´s t-test was performed for calculation of p-values for single parameters fresh weight excluding the roots, the length of the longest leaf, and the height of the vegetation point.
The discoloration of the hypocotyl was determined by observation of cross sections. To test for fungal outgrowth from stems of infected plants stems were harvested and surface sterilized by washing with 70% (v/v) ethanol for 8 min, followed by 6% (v/v) sodium hypochloride solution for 8 min and subsequent washing with distilled sterile water, twice. The stem ends were removed and slices of the middle part were placed on PDM plates supplemented with chloramphenicol (100 µg/ml). Plates were incubated at 25 °C for 7 d and fungal growth was documented.

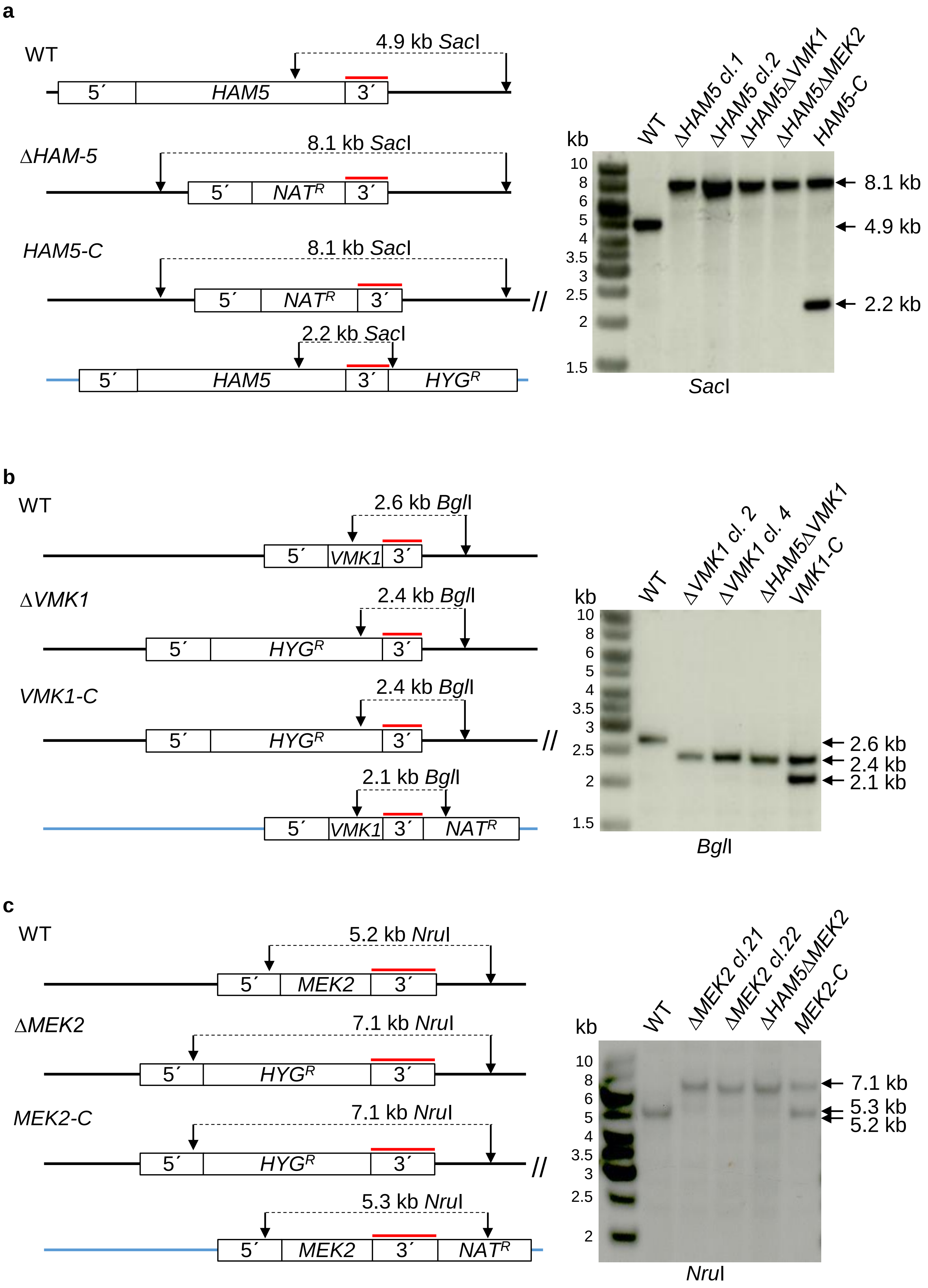


**Supporting Information Figure S1**

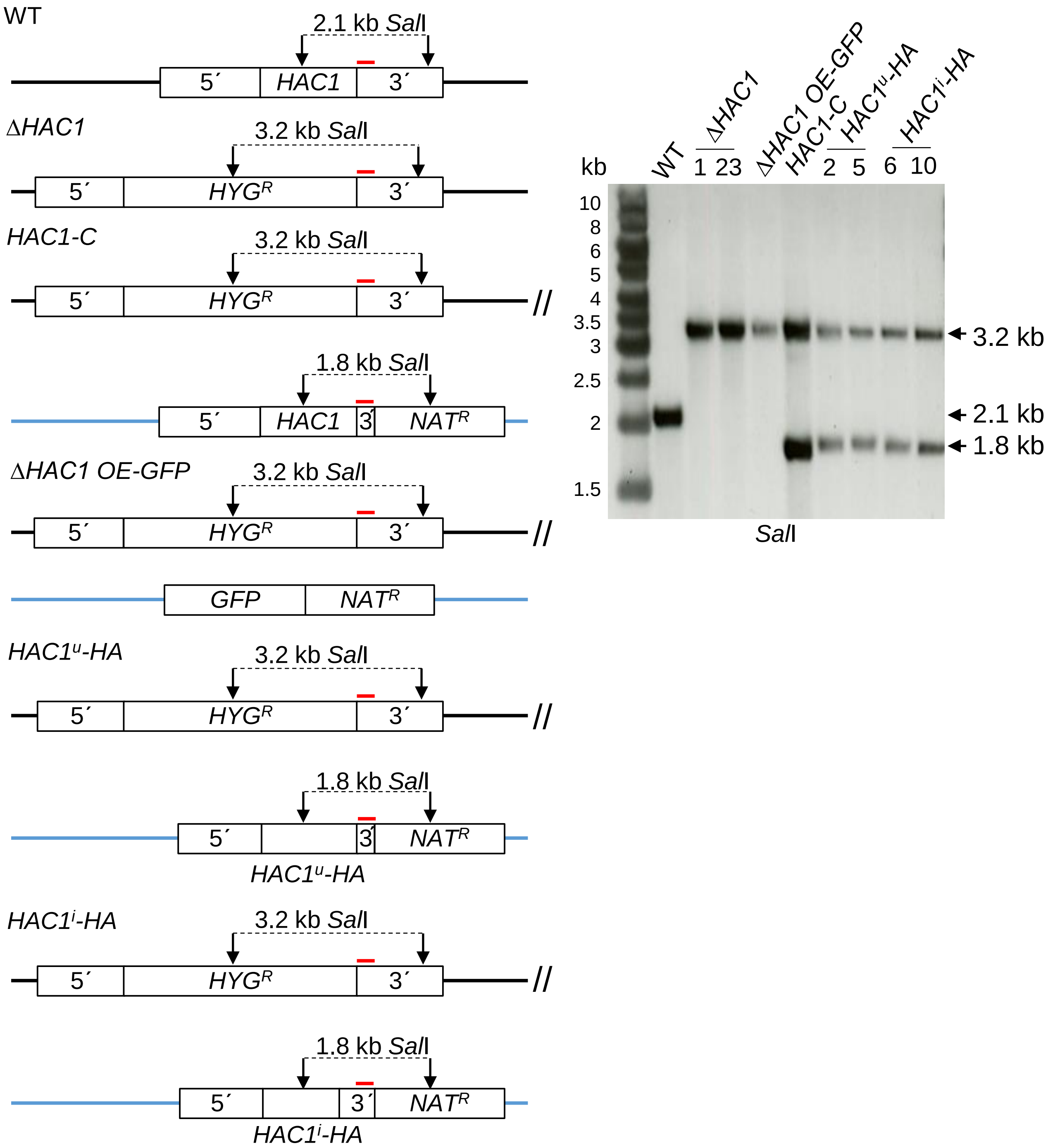


**Supporting Information Figure S2**

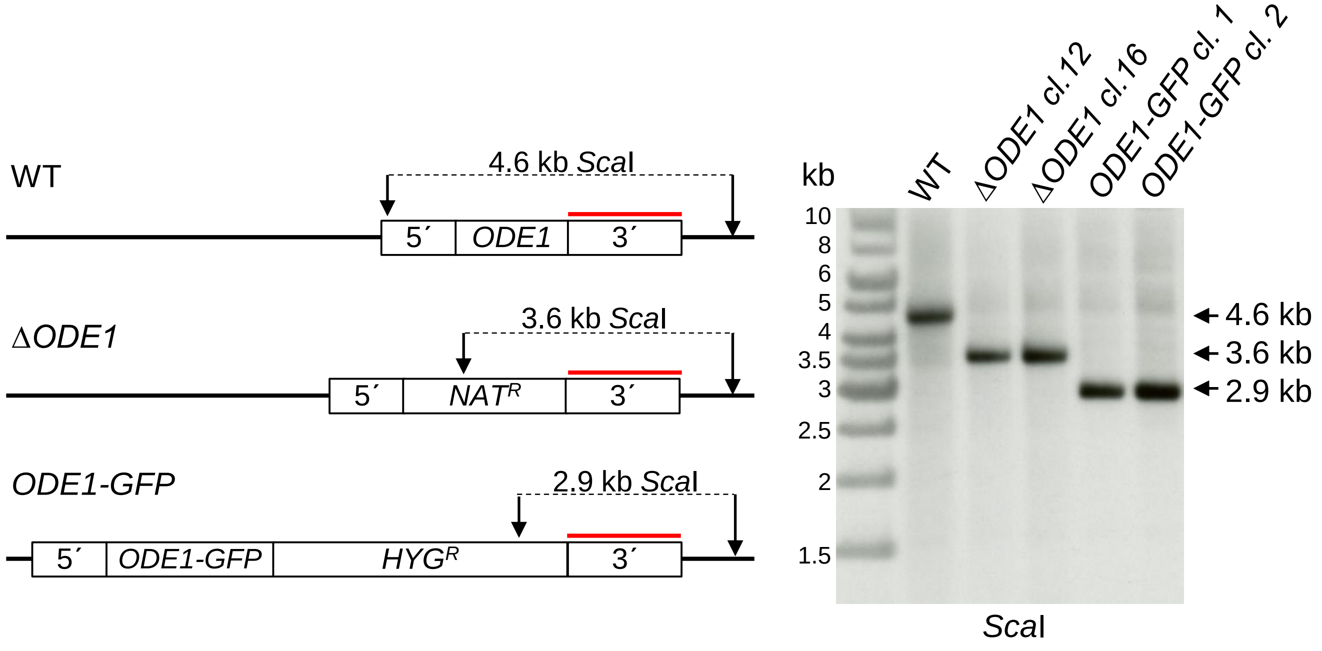


**Supporting Information Figure S3**

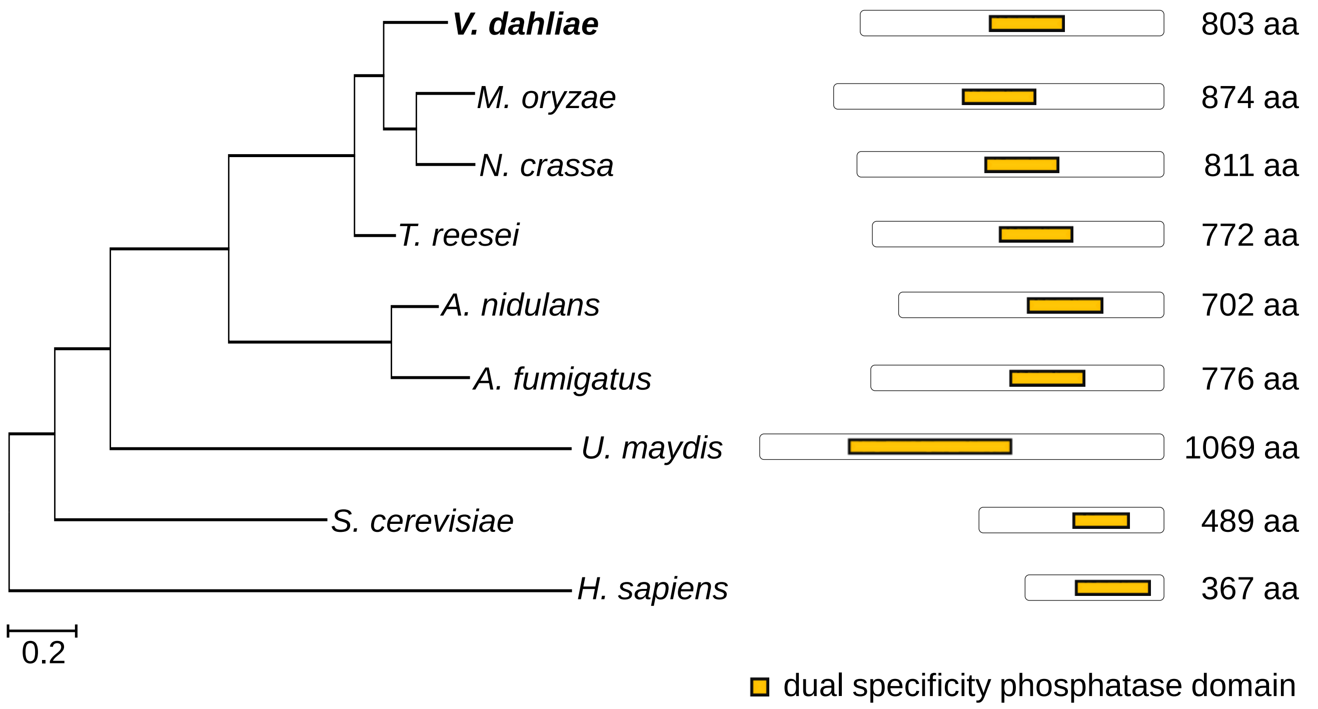


**Supporting Information Figure S4**
